# *Fusarium graminearum* DICER-like-dependent sRNAs are required for the suppression of host immune genes and full virulence

**DOI:** 10.1101/2021.05.17.444440

**Authors:** Bernhard Werner, Aline Koch, Ena Šečić, Jonas Engelhardt, Lukas Jelonek, Jens Steinbrenner, Karl-Heinz Kogel

**Affiliations:** Institute of Phytopathology, Centre for BioSystems, Land Use and Nutrition, Justus Liebig University, Heinrich-Buff-Ring 26-32, D-35392, Giessen, Germany; Institute for Phytomedicine, University of Hohenheim, Otto-Sander-Straße 5, D-70599, Stuttgart, Germany; Institute of Bioinformatics and Systems Biology, Justus Liebig University, Heinrich-Buff-Ring 58, D-35392, Giessen, Germany

**Keywords:** RNAi, sRNA, Fusarium graminearum, DCL, Barley, Hordeum vulgare, Brachypodium distachyon, SERK, EOL1, BAK1

## Abstract

In filamentous fungi, gene silencing by RNA interference (RNAi) shapes many biological processes, including pathogenicity. Recently, fungal small RNAs (sRNAs) have been shown to act as effectors that disrupt gene activity in interacting plant hosts, thereby undermining their defence responses. We show here that the devastating mycotoxin-producing ascomycete *Fusarium graminearum* (*Fg*) utilizes DICER-like (DCL)-dependent sRNAs to target defence genes in two Poaceae hosts, barley (*Hordeum vulgare Hv*) and *Brachypodium distachyon* (*Bd*). We identified 104 *Fg*-sRNAs with sequence homology to host genes that were repressed during interactions of *Fg* and *Hv*, while they accumulated in plants infected by the DCL double knock-out (dKO) mutant PH1-*dcl1/2*. The strength of target gene expression correlated with the abundance of the corresponding *Fg*-sRNA. Specifically, the abundance of three tRNA-derived fragments (tRFs) targeting immunity-related *Ethylene overproducer 1-like 1* (*HvEOL1)* and three Poaceae orthologues of *Arabidopsis thaliana BRI1-associated receptor kinase 1* (*HvBAK1, HvSERK2* and *BdSERK2*) was dependent on fungal DCL. Additionally, RNA-ligase-mediated Rapid Amplification of cDNA Ends (RLM-RACE) identified infection-specific degradation products for the three barley gene transcripts, consistent with the possibility that tRFs contribute to fungal virulence via targeted gene silencing.

**Significance Statement:** *Fusarium graminearum* is one of the most devastating fungal pathogens in cereals, while understanding the mechanisms of fungal pathogenesis is a prerequisite for developing efficient and environmentally friendly crop protection strategies. We show exploratory data suggesting that fungal small RNAs play a critical role in Fusarium virulence by suppressing plant immunity.

## Introduction

RNA interference (RNAi) is a biological process in which small RNA (sRNA) molecules mediate gene silencing at the transcriptional or post-transcriptional level. In agriculture, RNAi-mediated silencing strategies have the potential to protect crops from pests and microbial pathogens (Koch & Kogel 2014; Guo et al. 2016; Liu et al. 2020; Šečić & Kogel 2021; Koch & Wassenegger 2021). Expression of non-coding double-stranded (ds) RNA targeting essential genes in a pest, a pathogen or a virus can render host plants more resistant by a process known as host-induced gene silencing (HIGS) (Rosa et al. 2018; Cai et al. 2018a; Gaffar & Koch 2019; Niehl & Heinlein 2019). Alternatively, plants can be protected by foliar application of dsRNA to plants (Koch et al. 2016; Wang et al. 2016a; Konakalla et al. 2016; Mitter et al. 2017; Kaldis et al. 2018; McLoughin et al. 2018; Sang et al. 2020). While these RNAi-based crop protection strategies are proving to be efficient and agronomically practical in the control of insects (Head et al. 2017) and viruses (Niehl et al. 2018), many questions remain unanswered with regard to the control of fungi.

The blueprint for using RNA to fight disease comes from nature (Cai et al. 2018a). During infection of *Arabidopsis thaliana* (*At*), the necrotrophic ascomycete *Botrytis cinerea* (*Bc*) secretes DICER-like (DCL)-dependent sRNAs that are taken up into plant cells to interact with the Arabidopsis ARGONAUTE protein *At*AGO1 and initiate silencing of plant immune genes (Weiberg et al. 2013; Cai et al. 2018b). For instance, sRNA *Bc*-siR3.2 targets mitogen-activated protein kinases, including *MPK2* and *MPK1* in *At*, and *MAPKKK4* in tomato (*Solanum lycopersicum*), while *Bc*-siR37 targets several immune-related transcription factors including *WRKY7*, *PMR6* and *FEI2* (Wang et al. 2017b). Likewise, the oomycete *Hyaloperonospora arabidopsidis* produces 133 AGO1-bound sRNAs, which are crucial for virulence (Dunker et al. 2020), and microRNA-like RNA1 (*Pst*-milR1) from the yellow rust causing biotrophic basidiomycete *Puccinia striiformis* f.sp. *tritici* (*Pst*) reduced expression of the defence gene *Pathogenesis-related* 2 (*PR2*) in wheat (*Triticum aestivum*) (Wang et al 2017a). Notably, when comparing sRNA in the leaf rust fungus *Puccinia triticina* (*Pt*), 38 *Pt*-sRNAs were homologous to sRNAs previously identified in *Pst* (Dubey et al. 2019; Mueth et al. 2015), hinting to the possibility that sRNA effectors are conserved among related fungal species as it is known for plant miRNAs (Reinhart et al. 2002, Jones-Rhoades 2012). One group of conserved sRNAs with putative effector function are transfer RNA (tRNA)-derived fragments (tRFs). Bacterial tRFs play a role in the symbiotic interaction between soybean (*Glycine max*) and its nitrogen fixing symbiont *Bradyrhizobium japonicum* during root nodulation (Ren et al. 2019). Similarly, the protozoan pathogen *Trypanosoma cruzi* secretes tRF-containing microvesicles resulting in gene expression changes in mammalian host cells (Garcia-Silva et al. 2014).

Fungal species of the genus *Fusarium* belong to the most devastating pathogens of cereals causing Fusarium head blight and crown rot (Dean et al., 2012), and contaminate the grain with mycotoxins such as the B group trichothecenes deoxynivalenol (DON), nivalenol (NIV), and their acetylated derivatives (3A-DON, 15A-DON, and 4A-NIV) (Desjardins et al., 1993; Jansen et al., 2005; Ilgen et al., 2009). Viability, aggressiveness, and virulence of Fusaria are under control of the RNAi machinery (Kim et al., 2015; Son et al., 2017; Gaffar et al. 2019).

To test the possibility of *Fg* producing sRNAs that exert effector function and promote pathogenesis, we predicted Fg-sRNA targets in two Poaceae hosts, *Hordeum vulgare* (*Hv*) and *Brachypodium distachyon* (*Bd*). Among the many predicted plant targets of fungal sRNA, three fungal tRFs had sequence similarity to *BRI1-associated receptor kinase 1* (*BAK1*) homologs and *EOL1* (*Ethylene overproducer 1-like 1*) in *Hv* and *Bd*. Upon infection with the wild type *Fg* strain, transcripts of genes were strongly reduced, while in contrast they were increased upon infection with *Fg* strains compromised for DCL activity. Degradation products of target mRNAs were detected by RNA-ligase-mediated Rapid Amplification of cDNA Ends (RLM-RACE), supporting the possibility that DCL-dependent sRNAs play a critical role in the interaction of *Fg* with cereal hosts.

## Results

### Fusarium graminearum DCL mutants are less virulent on barley and Brachypodium leaves

The Fusarium mutant IFA65-*dcl1* is partially impaired in infecting wheat ears and causing Fusarium head blight (Gaffar et al. 2019). We extended this earlier study to examine the effects of impaired DCL activity on the plant defence response. To this end, two to three-week-old detached second leaves of barley cv. Golden Promise (GP) were drop-inoculated with 3 µl of a solution containing 150,000 conidia per ml of *Fg* isolate PH1 or the double knock-out (dKO) mutant PH1-*dcl1/2*. At five days post inoculation (dpi), the dKO mutant produced significantly smaller necrotic lesions (30%; median (MED) (27%); interquartile range (IQR) (47%) Wilcoxon rank sum test, p=0.007) than the wild type (wt) strain, confirming that DCL activity is required for full *Fg* virulence (Fig. 1A).

**Fig. 1:**
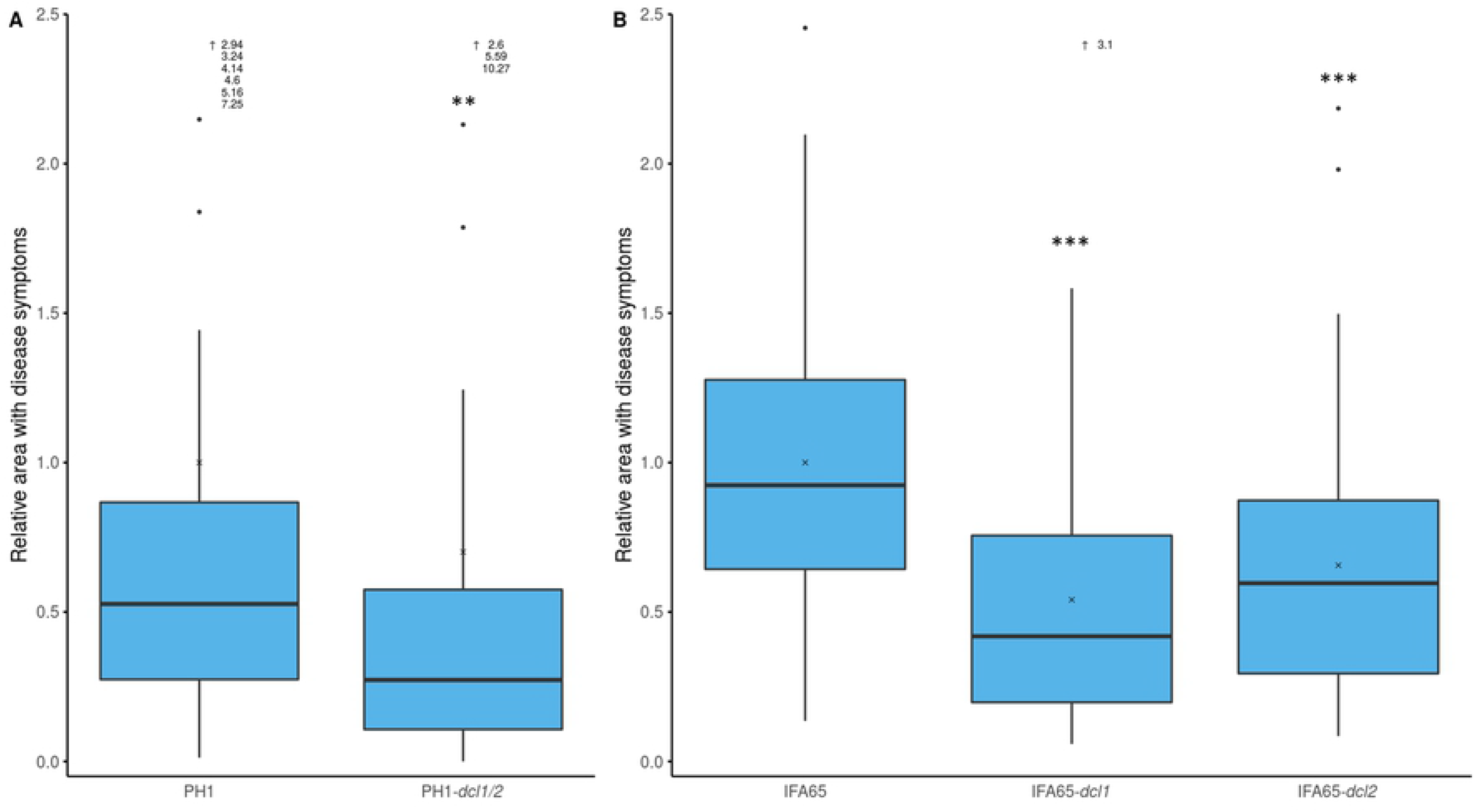
Virulence of Fusarium graminearum strains PH1 and IFA65 DCL single and dKO mutants on barley and Brachypodium. A: Relative infected area on leaves of barley cv. Golden Promise (GP) at 5 dpi. Detached leaves were inoculated with 3 µl of a solution containing 150,000 conidia per mL. The area with leaf necrosis was measured with the free image analysis software package PlantCV. Boxplots represent the median and quartiles of three independent biological experiments (n=56). (Wilcoxon Rank Sum Test, P=7.1*10^-3^, x=mean) B: Relative infected leaf area on leaves of *Brachypodium distachyon* Bd21-3 at 5 dpi. Detached leaves were inoculated with 10 µl of a solution containing 10,000 conidia per mL. The area with leaf necrosis was measured with ImageJ. Boxplots represent the median and quartiles of nine independent biological experiments (n=63). (Pairwise Wilcoxon Rank Sum Test, Bonferroni corrected, P*_dcl1_*=4.9*10^-8^, P*_dcl2_*=2.6*10^-4^, x=mean) Outliers (>2.5) are not shown but indicated as written values next to the upward arrow (〈).

Next, we determined the virulence of *DCL* mutants on *Brachypodium distachyon* Bd21-3. Flag leaves of three-week-old plants were inoculated with 10 µl (10,000 conidia ml^-1^) of fungal inoculum. Single mutants IFA65-*dcl1* and IFA65-*dcl2* produced significantly smaller lesions than the wt (IFA65-*dcl1*, 54%; MED (42%); IQR (56%) and IFA65-*dcl2*, 66%; MED (60%); IQR (58%); pairwise Wilcoxon rank sum test; Bonferroni corrected; p<0.005) (Fig. 1B). These results substantiate the earlier findings (Gaffar et al. 2019) that fungal DCL activity is required for Fusarium virulence on graminaceous plants.

### Selection of sRNAs with sequence homology to plant genes

We looked for interaction-related fungal sRNAs that potentially could interfere with plant gene expression by sequence-specific silencing. To this end, a previously published sRNA sequencing data set of *Fg* sRNAs from an axenic IFA65 culture (Koch et al. 2016) was analysed for sRNAs with sequence complementarity to barley genes. In order to identify a wide range of potential targets, we applied only two selection criteria, namely *i.* size (21-24 nt) and *ii.* A minimal number of reads (at least 400 reads in the dataset). From a total of 35,997,924 raw reads, 5,462,596 (comprising 589,943 unique sequences) had a length of 21-24 nt. From the unique sequences, 1,987 had at least 400 reads. Since the IFA65 genome has not been sequenced, we used the published genome information of *Fg* strain PH1 (genome assembly ASM24013v3 from International Gibberella zeae Genomics Consortium: GCA_000240135.3) for further analysis. The majority of the 1,987 unique sRNAs mapped to rRNA (64.4%) and intergenic regions (21.6%), while 3.7% and 2.4% mapped to protein coding genes and tRNAs, respectively, and 7.8% did not perfectly match the reference genome (Fig. S1). According to the TAPIR algorithm, the 1,987 sequences overall matched mRNAs of 2,492 genes (Hordeum vulgare IBSC PGSB v2 reference genome; Mascher et al., 2017) sufficiently close according to the refined target prediction criteria suggested by Srivastava et al. (2014). GO-enrichment analysis revealed an enrichment in functions of nucleotide binding, motor activity and kinase activity and processes such as transport and localization (Fig. S2). Most of the 14,156 transcripts of the 2,492 target genes, which we nominated as potential sRNA targets, showed partially homologous sequences to more than one sRNA accounting for a total of 17,275 unique pairs of potential target gene -sRNA combinations. Target prediction results are presented with only one transcript (splice variant) for every combination (Tab. S2). Of note, merely 101 out of the 1,987 sRNAs had no predicted target among the total number of 248,391 plant mRNAs in the IBSC_PGSB_v2 annotation.

### Barley immune genes accumulate to higher levels in PH1-dcl1/2-infected leaves

From the set of 2,492 barley genes with partial sequence homology to *Fg* sRNAs, we selected 16 genes for further analysis, based on an educated guess that they are potentially involved in biotic stress reactions during plant-fungal interaction (Tab. 1). When tested with RT-qPCR, we found eight genes significantly higher expressed (Student’s *t*-test, paired, *p<0.1, **p<0.05, ***p<0.01) in leaves infected with PH1-*dcl1/2* vs. PH1 (Fig. 2). Among these genes are three that encode proteins involved in the regulation of either ethylene (ET) (Ethylene overproducer 1-like 1, *Hv*EOL1) or auxin responses (Auxin response transcription factors *Hv*ARF10 and *Hv*ARF19) and three kinases, of which Somatic embryogenesis receptor-like kinase 2 (*Hv*SERK2) and BRI1-associated receptor kinase 1 (*Hv*BAK1) are likely involved in recognition of microbe-associated molecular patterns (MAMPs). Moreover, genes encoding the plastid kinase 2-Phosphoglycolate phosphatase 2 (*Hv*PGLP2), Resurrection 1 (*Hv*RST1, with a rather elusive function in cuticle formation and embryo development), and the histone-lysine N-methyltransferase Su(var)3-9-related protein 5 (*Hv*SUVR5, involved in transcriptional gene silencing) were also strongly expressed.

**Table 1:**
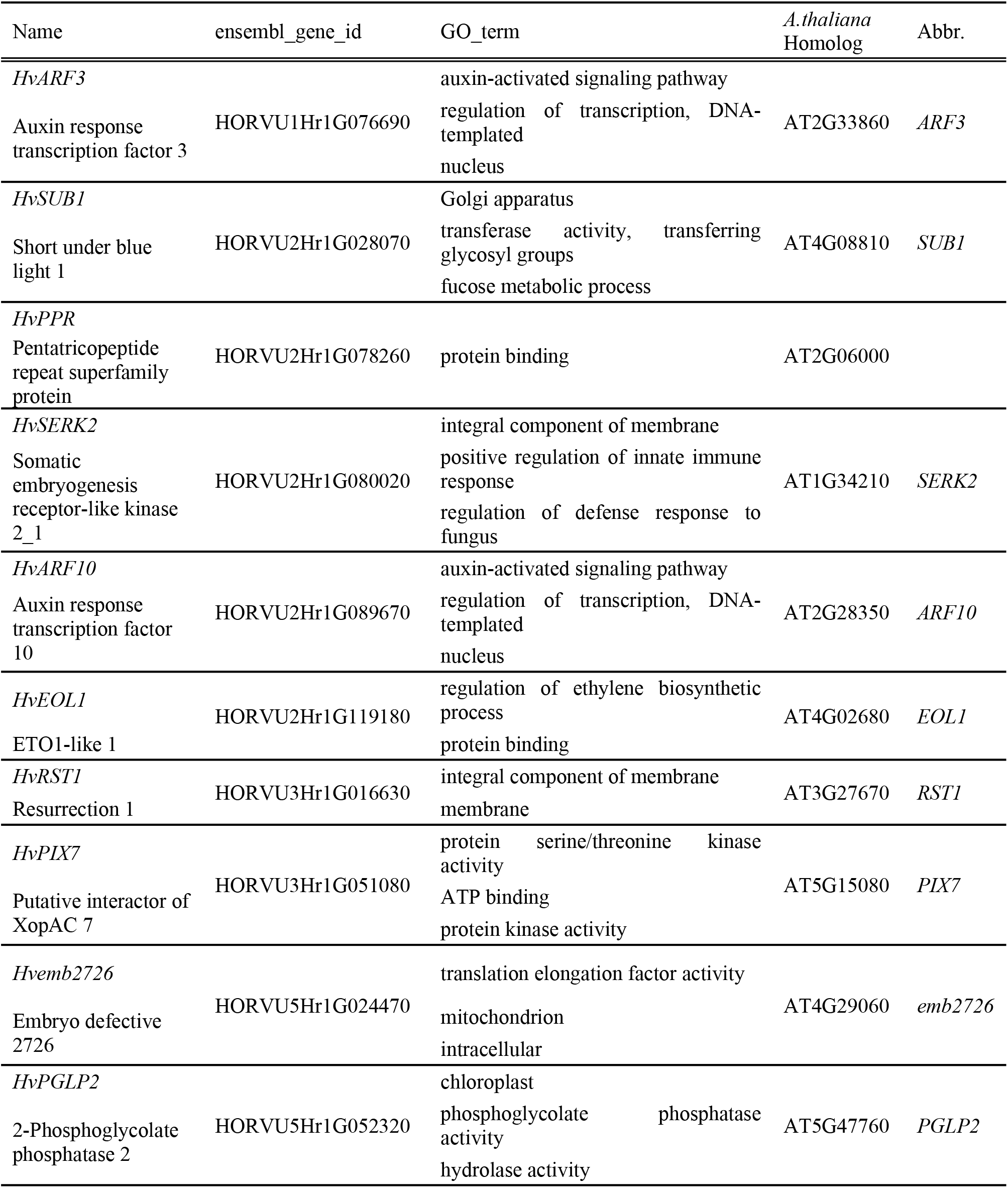
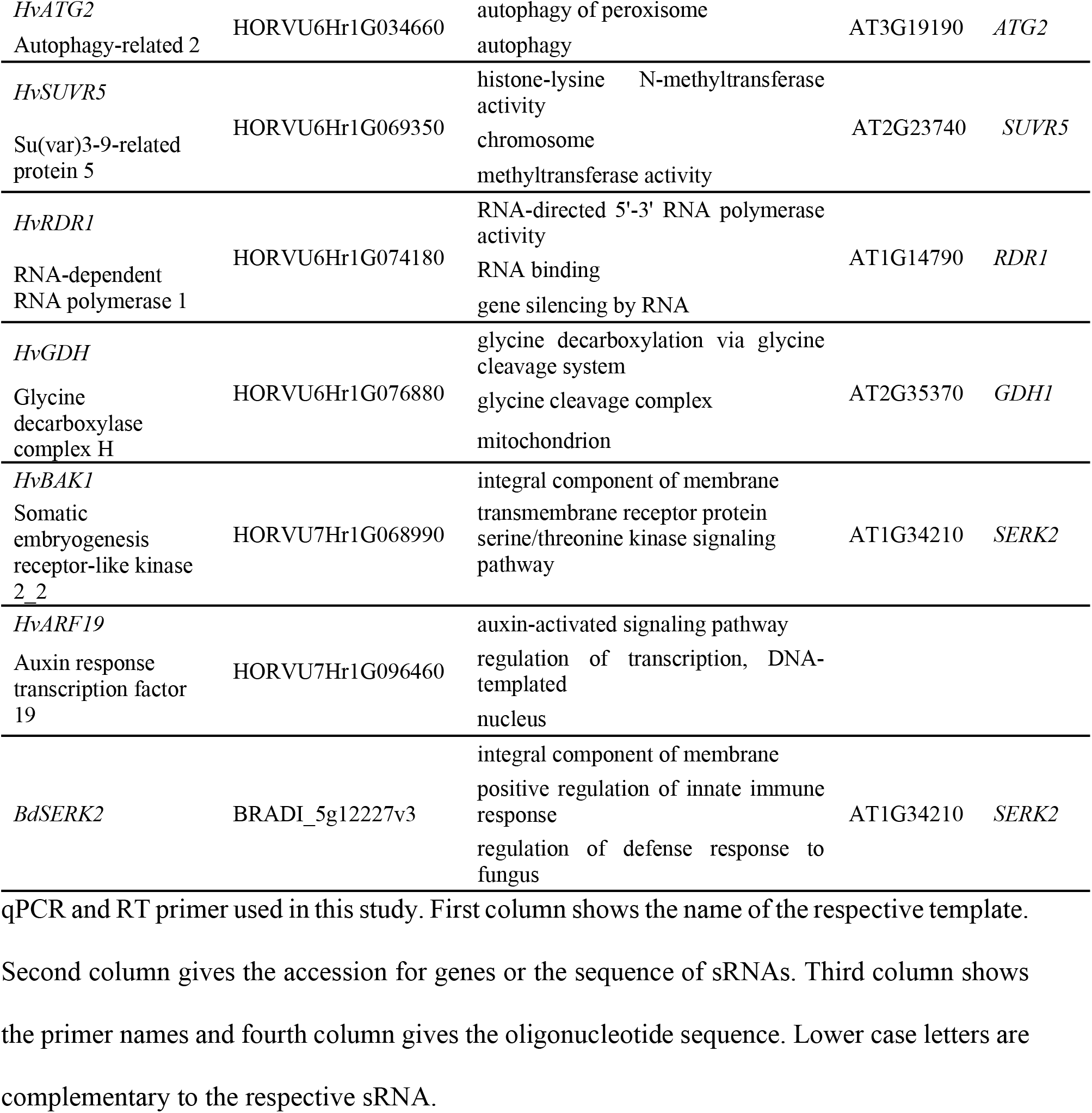
Selected GO-terms of tested genes and closest homologs in A. thaliana.

**Fig. 2:**
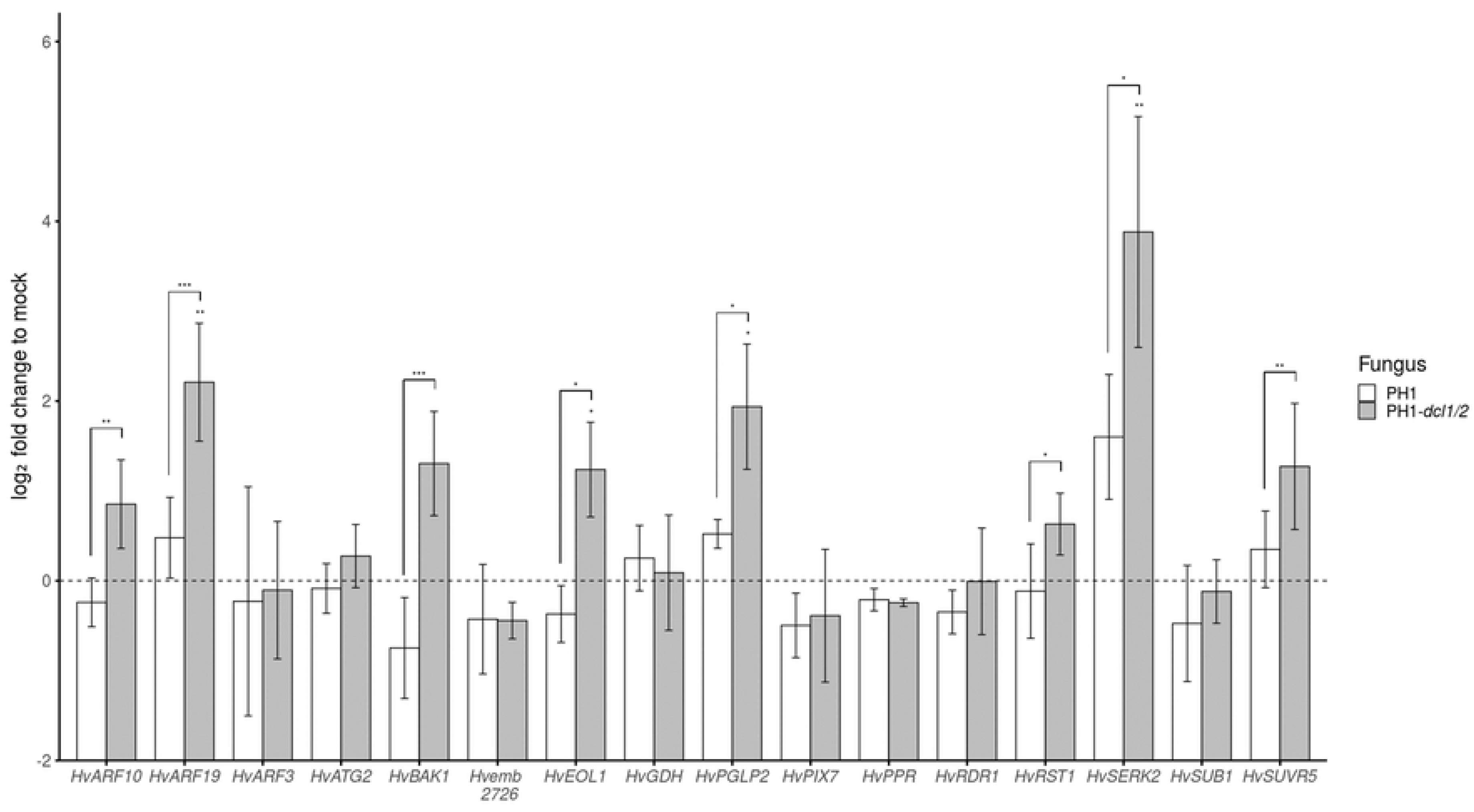
Relative expression (log2 fold) of potential barley target genes for fungal sRNAs in leaves infected with Fusarium graminearum wt strain PH1 vs. PH1-dcl1/2. Expression was normalized against barley *Ubiquitin* (*HvUBQ*) and subsequently against the Δct of the uninfected control (mock treatment). Bars represent the mean±SE of three independent biological replicates. Significant differences were calculated for the expression of a respective gene in PH1 vs. PH1-*dcl1/2*-infected samples and PH1 vs. controls. The dotted line shows the expression level of mock treatment. (Student‘s *t*-test, (paired) one sided, *P<0.1, **P<0.05, ***P<0.01)

### HvEOL1 transcripts also accumulate to higher levels upon DCL knock-down via spray induced gene silencing (SIGS)

We selected *HvEOL1* (*HORVU2Hr1G119180*), which is a homologue of *At Ethylene overproducer1* (*AtETO1*; *AT4GO2680.1*), for further analysis. The alignment of the respective protein sequences of *Hv*EOL1 and *At*ETO1 is shown in Fig. S3. *At*ETO1 negatively regulates ethylene synthesis in *At* by ubiquitination of type-2 1-Aminocyclopropane-1-carboxylate synthases (ACSs), which produce the direct precursor of ET (Christians et al. 2009) (Fig. S4). Upon inoculation with PH1, *HvEOL1* expression was reduced by 23% as compared to non-inoculated barley leaves. In contrast, *HvEOL1* was strongly expressed in PH1-*dcl1/2*-infected leaves well above the levels measured either in PH1- or mock-inoculated leaves. To further substantiate that *HvEOL1* expression is under the control of fungal DCL activity, we used a SIGS strategy (Koch et al. 2016) to partially inactivate DCL function in *Fg*. Two-week-old detached leaves were sprayed with 20 ng µl^-1^ of dsRNA-*dcl1/2*, a 1,782 nt long dsRNA derived from the sequences of IFA65-*DCL1* and IFA65-*DCL2* (Fig. S5A,B). 48 h later, leaves were drop inoculated with conidia and harvested at 5 dpi. Consistent with the expectation that exogenous dsRNA-*dcl1/2* mediates silencing of their *DCL* gene targets, RT-qPCR analysis confirmed that the transcript levels of IFA65*-DCL1* and IFA65-*DCL2* were reduced to 22% and 42%, respectively, as compared with the Tris-EDTA (TE) buffer control (Fig. 3A). In accordance with the results obtained with strain PH1, *HvEOL1* was also significantly (p=0.029, Student’s *t*-test (Δct), one sided, paired) downregulated in response to IFA65 infection compared to mock controls treated with 0.02% Tween20 (Fig. 3B). In contrast, however, when leaves were sprayed with dsRNA-*dcl1/2* prior to inoculation with IFA65, *HvEOL1* transcripts strongly accumulated in comparison with the inoculated leaves sprayed with TE buffer (Fig. 3C).

**Fig. 3:**
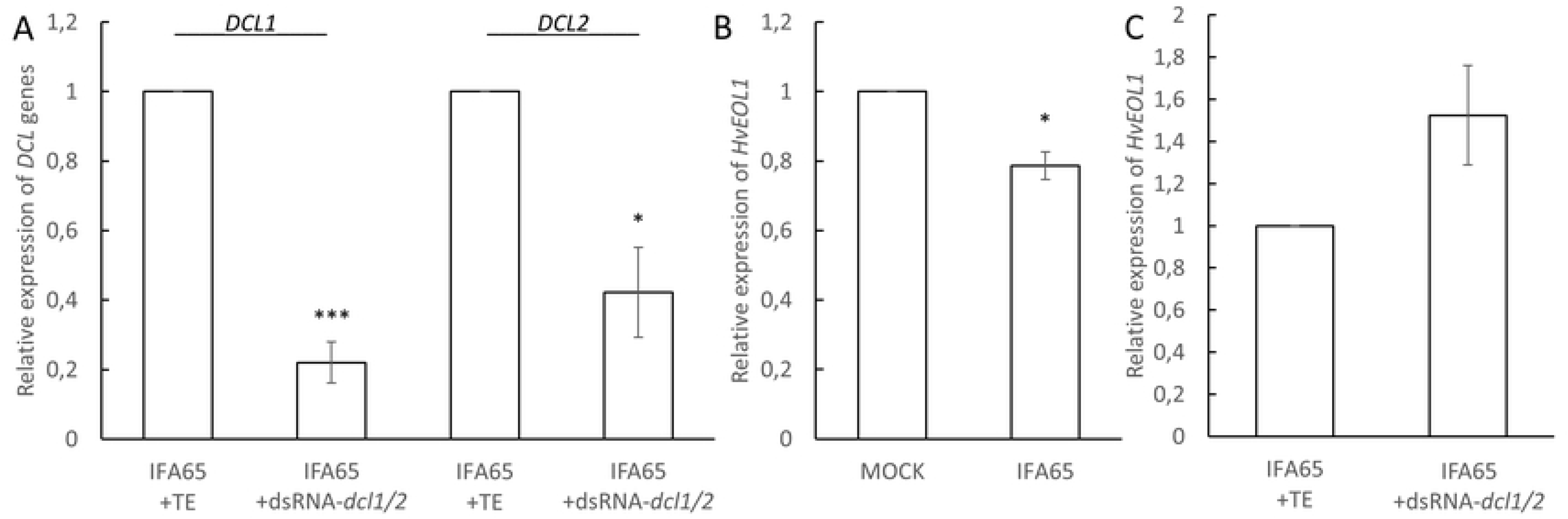
Relative expression of HvEOL1 in response to inoculation of barley leaves with Fusarium graminearum. A, Relative expression of *FgDCL1* and *FgDCL2* on detached barley cv. Golden Promise leaves at 5 dpi in wt strain IFA65 and 7 days post spray application of the 1,782 nt long dsRNA construct dsRNA-*dcl1/2* vs. TE buffer. **B**, Relative expression of *HvEOL1* at 5 dpi with IFA65 vs. mock control. **C**, Relative expression of *HvEOL1* 5 dpi with wt strain IFA65 and 7 days post spray application of dsRNA-*dcl1/2* vs. TE buffer. Gene expression was first normalized against the reference gene *HvUBQ (HORVU1Hr1G023660)* and subsequently against the Δct of the respective control for B (mock = 0.002% Tween20) and for A,C (IFA65 / TE). Bars represent the mean ± SE of three (B) and four (A, C) independent biological replicates. (Student‘s *t*-test, *P<0.05, ***P<0,005)

### Fungal sRNAs targeting HvBAK1, HvEOL1, and HvSERK2 mRNAs are less abundant in PH1-dcl1/2 vs. PH1

To detect the abundance of specific *Fg*-sRNAs, originally identified by sequencing of axenic IFA65 mycelium, in PH1-infected plant tissue, we performed reverse transcription stem-loop qPCR (Chen et al. 2005). From the above defined pool of 1,987 *Fg*-sRNAs (axenic, 21-24 nt length, >=400 reads) 22 unique sRNAs matched partial sequences of *HvEOL1*, 10 matched *HvBAK1* and five matched *HvSERK2*. *Fg*-sRNA-1921 matched all three genes and *Fg*-sRNA-321 matched both *HvEOL1* and *HvBAK1* (Tab. 2; Tab. S2). These two sRNAs show high sequence similarities among each other. To identify their origin, they were aligned to the genomic sequence of strain PH1 (GCA_900044135.1). We found that they match the gene *Fg_CS3005_tRNA-Gly-GCC-1-9* encoding tRNA-Gly for the anticodon GCC. Of note, a larger cluster of 27 overlapping tRNA-derived fragments (tRFs) with more than 50 reads matching the tRNA-Gly gene sequence were detected (Fig. S6). To assess differential accumulation of tRFs from the *Fg_CS3005_tRNA-Gly-GCC-1-9* cluster in leaves infected with PH1 *vs.* PH1-*dcl1/2*, sRNAs were reverse transcribed using hairpin-priming followed by qPCR amplification (Chen et al. 2005). For this analysis, we chose *Fg*-sRNA-321, the most abundant tRF from this cluster, along with *Fg*-sRNA-1921, which targets all three GOIs and an additional tRF (*Fg*-sRNA-6717), which targets *HvEOL1* and *HvBAK1* (see Tab. 2) to assess the sensitivity of the assay. In the initial IFA65 dataset the *Fg*-sRNA-321 had a read count of 2,106, *Fg*-sRNA-1921 had 416 and *Fg*-sRNA-6717 had 86 from a total of more than 5 million reads (Fig. S7). This equals 386 reads per million (rpm) for *Fg*-sRNA-321, while in average unique reads had only 1.7 rpm. Using TAPIR (Bonnet et al. 2010), we also calculated the target score values for all three tRFs, which is a measure for the similarity between sRNA and target. A high value refers to more dissimilarities. Mismatches (MMs) increase the score by one point and G-U pairs by 0.5 points. These values are doubled if the respective MMs and G-U pairs are located between the second and 12^th^ nt of the sRNA (5’-3’) because a high similarity in the seed region of the sRNA is especially important for RNAi (Mallory et al. 2004). *Fg*-sRNA-321 has a score of 4.5 for *HvBAK1* and *HvEOL1*, *Fg*-sRNA-1921 has a score of 4, 3.5 and 6 for *HvBAK1*, *HvEOL1* and *HvSERK2*, respectively and *Fg*-sRNA-6717 has a score of 5.5 and 4.5 with *HvBAK1* and *HvEOL1*. In plants other than Arabidopsis, such as wheat and rice, a score cut off at 4 or 6 points lead to a precision of 82% or 62% and a recall of known interactions of 39% or 58% respectively according to Srivastava et al. (2014).

**Table 2:**
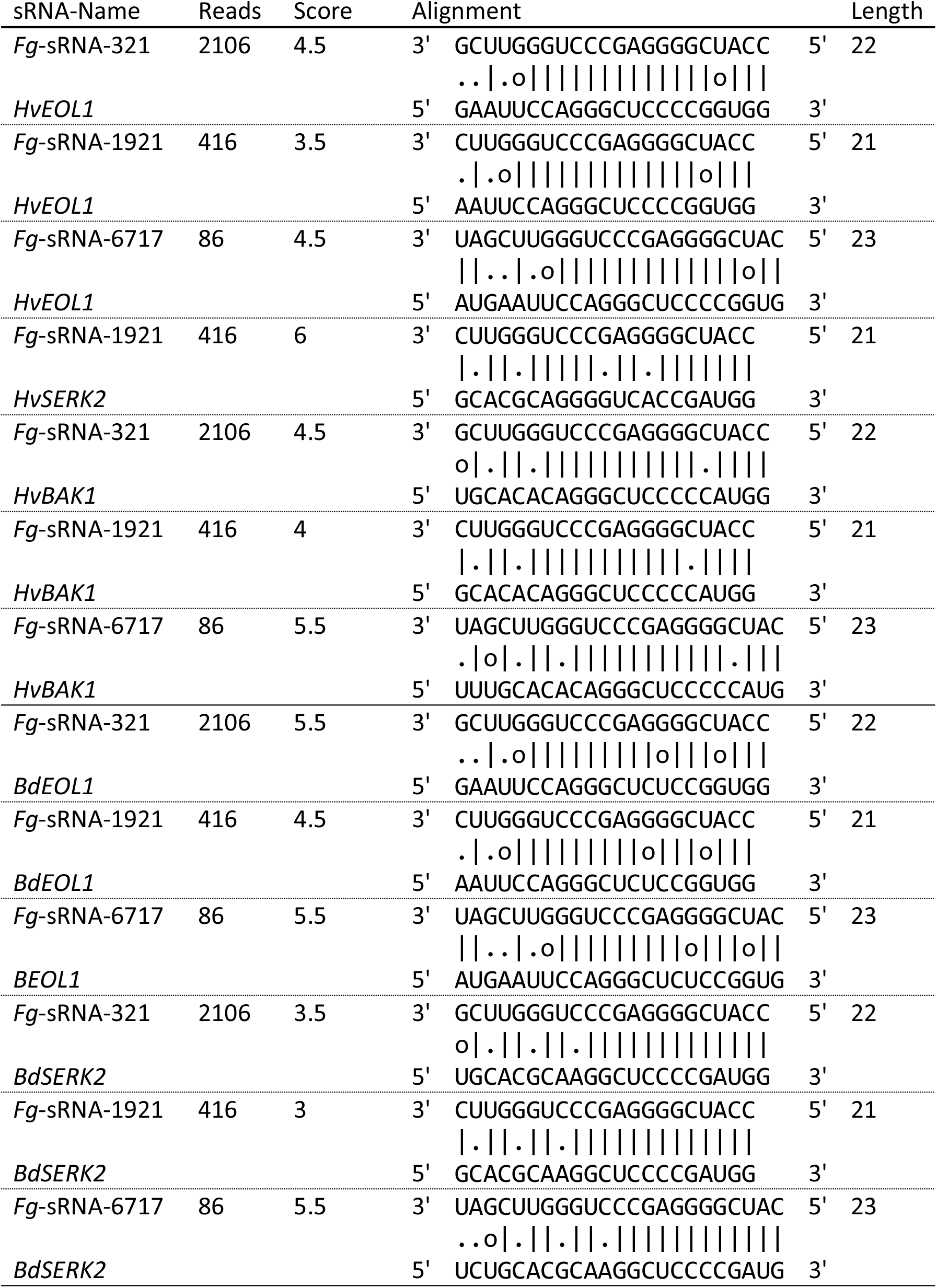
Target prediction results of Fg-sRNAs with more than 400 reads in IFA65 axenic culture.

All three fungal tRFs were detected in infected leaves, while they could not be found in uninfected leaves (Fig. 4). Significantly lower amounts of *Fg*-sRNA-1921 (59%), *Fg*-sRNA-321 (56%), and *Fg*-sRNA-6717 (60%) were detected in PH1-*dcl1/2 vs*. PH1-infected leaves (Fig. 4), showing that their biogenesis is DCL-dependent.

**Fig. 4:**
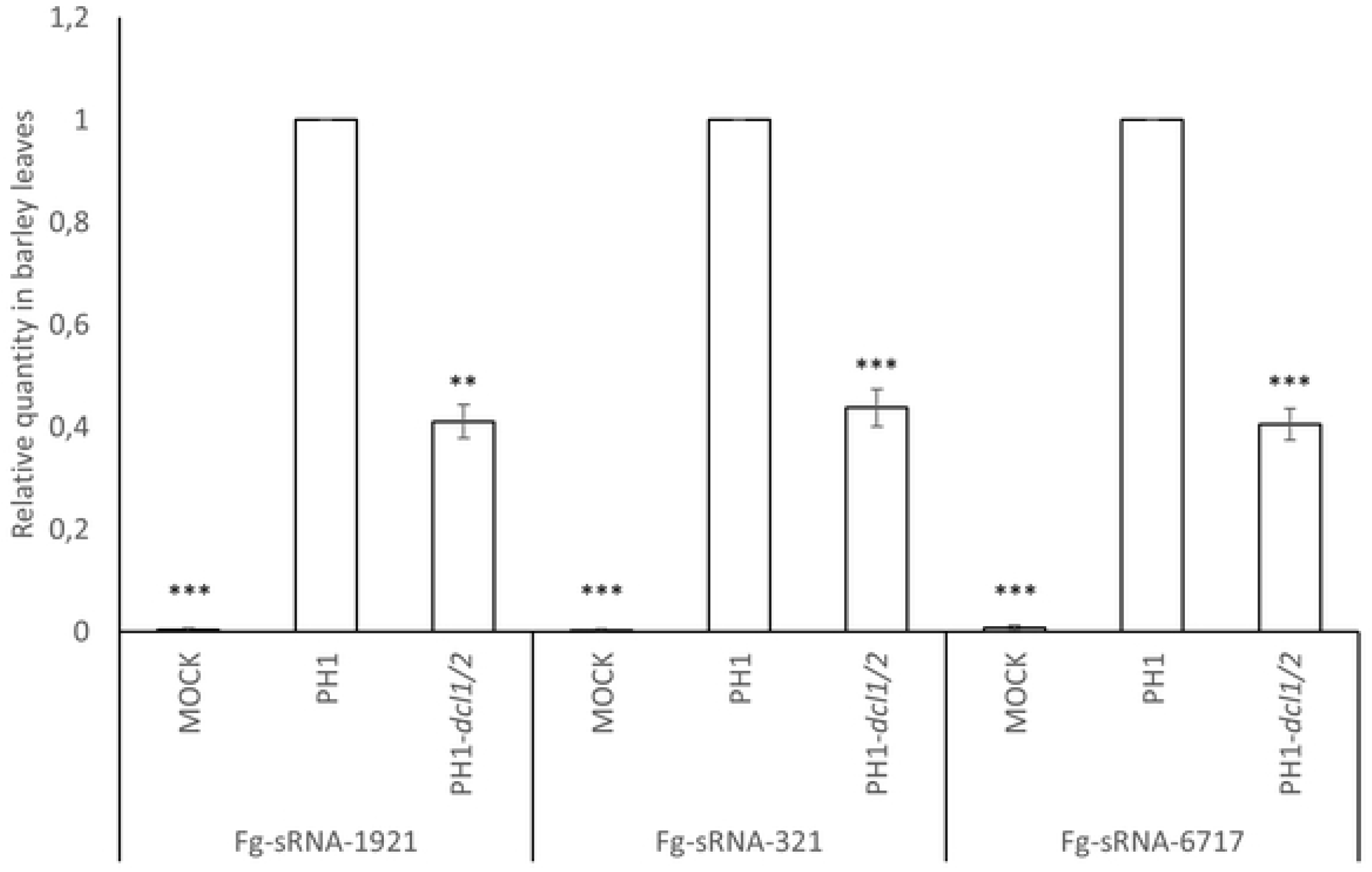
Relative amount of different fungal tRFs with homology to HvEOL1 mRNA. Relative amount of different fungal tRFs with homology to *HvEOL1* mRNA during infection of barley leaves with PH1 and PH1-*dcl1/2* normalized to fungal biomass and relative quantity of sRNAs normalized to wt PH1 measured by qPCR. *Fg*-sRNA-1921, *Fg*-sRNA-321 and *Fg*-sRNA-6717 quantity was normalized to *Hvu*-miR159 and *Hvu*-miR168 and fungal biomass as determined by *FgEF1α* expression was normalized to *HvUBQ*. Subsequently the amount of sRNAs was normalized with fungal biomass. The amount of sRNA in PH1-infected leaves was set to 1. Values and error bars represent the mean ± SE of three independent biological replicates. Significance was calculated via a one-sample *t*-test. (**P<0.01, ***P<0.005)

### Fg-sRNA-321 and Fg-sRNA-1921 also match SERK2 in Brachypodium distachyon Bd21-3

Next, we assessed the possibility that *Fg*-sRNA-321, *Fg*-sRNA-1921 and *Fg*-sRNA-6717 also have sequence homologies in *At* and the model grass *Bd*. Target prediction with the TAPIR algorithm using the optimised parameters for *At* (score=4; mfe=0.7), could not detect potential targets in *At* ecotype Col-0. In contrast, these three tRFs matched the sequence of *Brachypodium somatic embryogenesis receptor-like kinase 2* (*BdSERK2*) in Bd21-3 with a score of 3.5, 3 and 5.5, respectively (Tab. 2). We examined the expression pattern of *BdSERK2* in response to leaf infection: *BdSERK2* is relatively weakly expressed in uninfected plants and is not further suppressed after inoculation with PH1, whereas it strongly accumulated in PH1-*dcl1/2 vs.* PH1-infected Bd21-3 (Fig. 5). This finding further supports the possibility that the control of *SERK2* expression via RNAi pathways by *Fg* is evolutionary conserved in cereals.

**Fig. 5:**
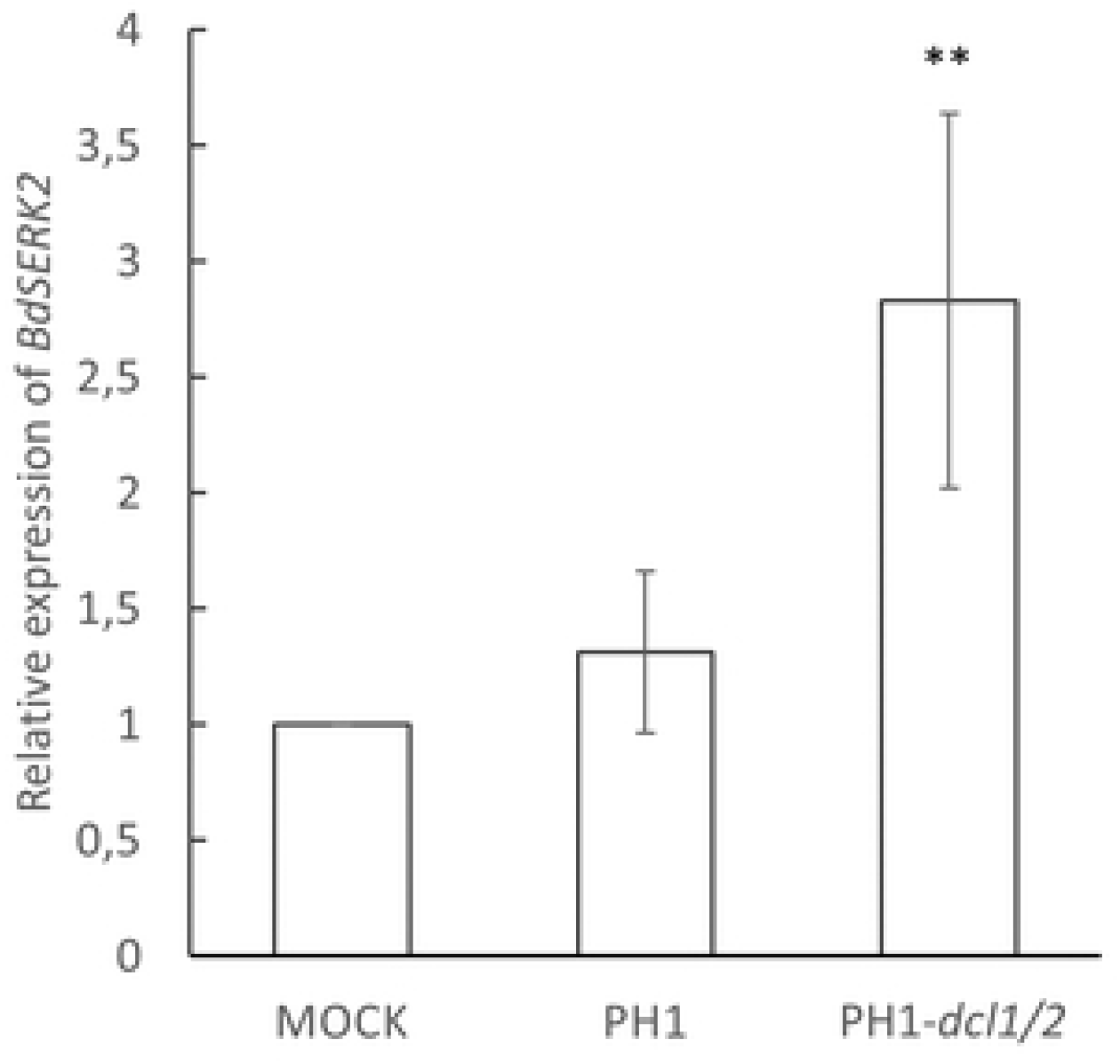
Relative expression of BdSERK2 in response to inoculation of Brachypodium distachyon leaves with Fusarium graminearum. Relative expression of *BdSERK2* in detached Bd21-3 leaves at 4 dpi with PH1 vs. PH1*-dcl1/2*. The gene expression was first normalized against the reference gene *BdUBI4* and subsequently against the Δct of the mock treated control. Values and error bars represent the mean ± SE of three independent biological replicates. (Student‘s t-test, paired, one sided, **P<0,01)

### RLM-RACE shows infection specific degradation products of HvBAK1, HvEOL1 and HvSERK2

We assessed the sRNA-mediated cleavage of *HvBAK1*, *HvEOL1*, and *HvSERK2* mRNAs, using a modified RNA-ligase-mediated Rapid Amplification of cDNA Ends (RLM-RACE) assay. Control samples were prepared both from uninfected tissue and from infected tissue without the reverse transcription step (no-RT control) and PCR products were visualized on an EtBr-Agarose gel. In these no-RT controls no amplification was visible.

For each gene more than one infection-specific product was amplified (blue and red arrows), which could not be amplified from the uninfected sample (Fig. 6D-F). We excised three bands (red arrows) of the expected size for a *Fg*-sRNA-1921 guided cleavage of *HvBAK1* (Fig. 6D) and one band for *HvEOL1* (Fig. 6E) and *HvSERK2* (Fig. 6F) and cloned them into the pGEM-T easy vector system. According to the IBSC_PGSB_v2 assembly, *HvBAK1* has splice variants, which could produce cleavage products of different lengths while for *HvSERK2* and *HvEOL1* there are no introns between sRNA target site and primer. From each band, five colonies were picked and for 23 of these extracted plasmids sequences were obtained. 16 sequences perfectly matched the reference genome, four with one MM and one with four MMs. Two sequences did not match the reference sufficiently enough to be aligned over the full length. The observed cleavage products are close to but do not match the canonical slice site between the 10^th^ and 11^th^ nt of *Fg*-sRNA-1921 and *Fg*-sRNA-321 (Fig. 6A-C).

**Fig. 6:**
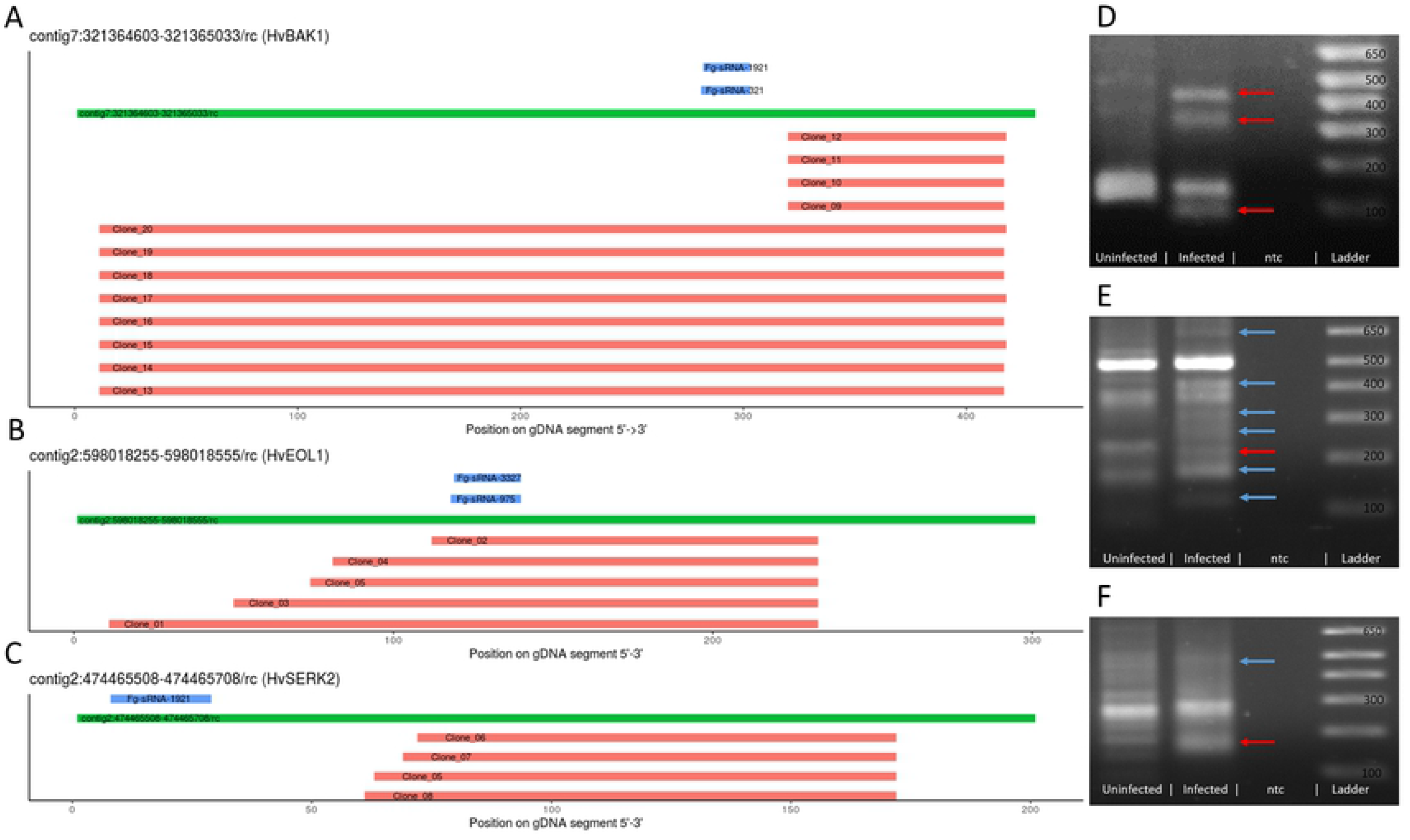
Analysis of potential target sites of Fg-sRNAs as determined by RLM-RACE products. A,B,C. Potential target sites of *Fg*-sRNA-321 and *Fg*-sRNA-1921 predicted by TAPIR (blue), genomic DNA (GPv1, GCA_902500625.1, A: contig7:321364603-321365033, B: contig2:598018255-598018555, C: contig2:474465508-474465708) of barley cv. Golden Promise (green), and the alignment of sequences derived from the RLM-RACE PCR (red) relative to the *Hv*-gDNA and *Fg*-sRNAs. **D,E,F** PCR-products of the second nested RLM-RACE-PCR visualized in an EtBr-Agarose gel. Red arrows indicate excised bands and blue arrows indicate infection specific products.

### Total sRNAs predicted to target a gene in barley are correlated with the de-repression strength

Not all potential targets of *Fg*-sRNAs are downregulated nor do all potential targets show a re-accumulation upon infection with PH1-*dcl1/2* (see Fig. 3). To address this bias we conducted a more focused target prediction exclusively for the 16 genes already tested by RT-qPCR. This allowed a much more thorough search, where targets for all sRNAs with at least two reads were predicted. From these 136,825 unique sRNAs (axenic, 21-24 nt length, >=2 reads) representing 4,997,312 reads of the total of 5,439,472 reads 21-24 nt in length, 5,052 have potential target sequences in the 16 mRNA sequences selected for further investigation in the *Hordeum vulgare* cv. GP assembly GCA_902500625. An additional filter step was employed to select for sRNAs with a maximum of one MM to the PH1 assemblies GCA_000240135.3 and GCA_900044135.1. Subsequently, sRNAs with up to one MM to *Fg*-rRNAs were removed leaving a total of 1,212 sRNAs with 1,311 potential sRNA-mRNA interactions representing 85,531 reads in the analysis.

To establish a correlation of the observed resurgence of potential target genes and targeting sRNAs, we analysed the *DCL*-dependent expression change using ΔΔΔct values. To compare the expression of a GOI in two samples, the difference between the ct-values for a reference gene and the GOI can be determined (Δct) and to calculate the expression difference between the control and treated sample the difference between the Δct values (ΔΔct) is calculated. We further defined the ΔΔΔct value as the difference between the ΔΔct values for a GOI in PH1 and PH1-*dcl1/2-*infected samples. From this follows a gene with a negative ΔΔΔct value shows a higher transcript accumulation during the infection with a fungal strain with compromised DCL function and the stronger the accumulation the lower this ΔΔΔct value is. We found a negative correlation between the ΔΔΔct value and the number of total sRNAs targeting a GOI (Fig. 7). This correlation becomes more significant if a lower score cut-off for the target prediction is chosen until the cut-off of four. The most significant correlation is for all predicted interactions with a score equal or below four with a p-value of 0.011 (t-test) (Fig. 7B). The p-value for a correlation with a cut-off of five (Fig. 7C) is 0.033 (t-test) and six (Fig. 7D) is 0.094 (t-test), while a score cut-off of 3 leads to a situation, where there are no predicted sRNA interactions for all genes except for three (Fig. 7A).

**Fig. 7:**
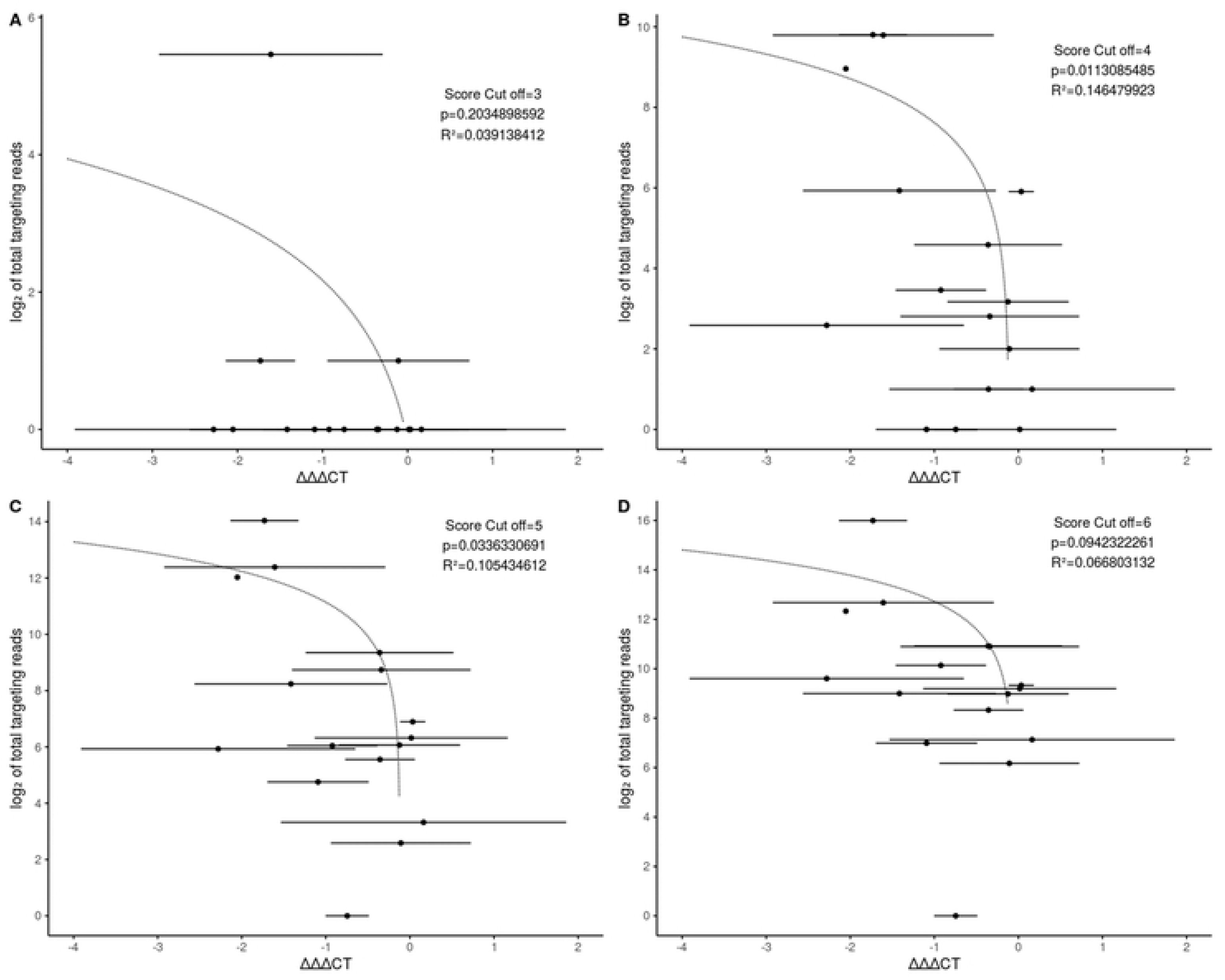
The degree of DCL-dependent gene silencing is correlated with the number of homologous fungal sRNAs. Each dot represents a predicted target gene of *Fg*-sRNAs. On the x-axis the ΔΔΔct-value is shown with bars representing SD. On the y-axis the log_2_ of the number of total sRNAs potentially targeting each gene are shown. The dotted line represents a linear regression model. P indicates the significance (t-test) of the model and the score cut-off indicates the score limit used during the target prediction. Plot A, B, C and D are the calculations for a score cut off of 3, 4, 5 and 6 respectively.

## Discussion

We show here that full virulence of the ascomycete fungus *Fusarium graminearum* on graminaceous leaves depends on the activity of fungal DCLs. The dKO mutant PH1-*dcl1/2* is less virulent on barley and the two single KO mutants IFA65-*dcl1* and IFA65-*dcl2* also are less virulent on *Brachypodium*. These results are consistent with our previous studies showing that knock-down or SIGS-mediated silencing of Fusarium *DCLs* and other components of the RNAi machinery reduced the virulence of the fungus on barley (Gaffar et al. 2019; Werner et al. 2020). DCL enzymes are key components of the fungal RNAi machinery required for the biogenesis of sRNAs directing silencing of sequence-complementary endogenous and foreign genes (Lax et al. 2020). The latter case involves DCL-dependent pathogen-derived sRNAs that target plant defense genes to increase virulence as shown for *Botrytis cinerea* (Weiberg et al. 2013; Wang et al. 2017b), *Puccinia striiformis* (Wang et al.2017a) and *Magnaporthe oryzae* (Zanini et al. 2021).

In the present work we found potential host target genes for fungal small RNAs (*Fg*-sRNAs) that were differentially regulated in response to plant infection with *Fg* wt *vs. Fg* DCL KO mutants, and the same effect was confirmed when DCLs were silenced by SIGS. This suggests a scenario in which impaired DCL function resulting in reduced fungal RNAi activity ultimately leads to de-repression of host target genes. Of note, target gene de-repression was also observed when the transcript was not significantly downregulated by the wt fungus during infection. This could be explained by a mutually neutralizing effect in which *Fg*-sRNAs continuously target genes for silencing, while concurrent plant immune responses are a trigger for up-regulation. Thus, one can speculate that these described effects reflect an abrogation of host-favouring upregulation by host immunity *vs*. pathogen-favouring downregulation by sRNA effectors.

We identified three tRFs predicted to target *BdSERK2*, *HvBAK1*, *HvEOL1* and *HvSERK2*. Unexpectedly, these tRFs are partially DCL-dependent, with a reduced abundance by more than 50% during infections with the dKO mutant PH1-*dcl1/2 vs.* wt PH1 based on fungal biomass. Current knowledge of tRFs in fungi and oomycetes suggests that their silencing activity is independent of DCL, as shown for *Sclerotinia sclerotiorum* (Lee Marzano et al. 2019) and *Phytophthora infestans*, where the production is partially dependent on AGO (Åsman et al. 2014). Furthermore, analysis of tRFs in Cryptococcus spp. revealed a RNAi-independent generation of tRFs and possible compensatory effects in an RNAi-deficient genotype (Streit et al. 2021). Interestingly however, the tRFs *Fg*-sRNA-321, *Fg*-sRNA-1921 and *Fg*-sRNA-6717 are neither 5’- or 3’ tRNA halves nor do they belong to any of the described tRF-1, tRF-2, tRF-4 or tRF-5 classes (Kumar et al. 2016a) applied by the tRFtarget database for animals, yeast (*Schizosaccharomyces pombe*) and the bacterium *Rhodobacter sphaeroides* (Li et al. 2021). When following the classification of the tsRBase used for all eukaryotic kingdoms and bacteria (Zuo et al. 2021), the three tRFs are classified as internal tRFs based on the origin within the mature tRNA. Interestingly, there are tRFs found in *Phytophthora sojae* starting in the anticodon loop and ending in the T loop of mature tRNAs (Wang et al. 2016b), which resembles the Fg-sRNA tRFs (Fig. S8).

We observed several infection-specific degradation products of the predicted host target genes *HvBAK1*, *HvEOL1* and *HvSERK2* for tRFs *Fg*-sRNA-321, *Fg*-sRNA-1921 and *Fg*-sRNA-6717. However, cleavage occurred outside the canonical miRNA cleavage site as defined by Mallory et al. (2004), though these genes are partially silenced during infection and silencing is apparently abolished upon infection with the DCL dKO mutant. While the canonical cleavage site for miRNA-directed cleavage in *At* is well defined, the tRF-directed cleavage observed by 5’ RACE of transposable elements in *At* (Martinez et al. 2017) and of defence-related genes during the infection of black pepper (*Piper nigrum*) with *Phytophthora capsici* (Asha & Soniya 2016) was found outside of the canonical cleavage site. Additionally, the identification of sRNA-directed cleavage sites in barley often leads to divergent findings. Ferdous et al. (2017) predicted ∼400 target genes for 11 presumably drought responsive miRNAs and found cleavage products for 15 targets overlapping the respective miRNAs alignment through degradome sequencing in the two barley cultivars Golden Promise (GP) and Pallas. From these confirmed targets, 13 were cleaved at the canonical 10^th^-11^th^ nt site, one was cleaved at 19^th^-20^th^ nt, and one at the 5^th^-6^th^ nt. Hackenberg et al. (2015) predicted 97 target genes of drought responsive miRNAs in GP and identified eight targets through degradome sequencing, which were all cleaved outside of the 10^th^-11^th^ nt site. Thus, both studies suggest the presence of non-canonical miRNA directed cleavage. Of note, both studies relied on the same degradome sequencing dataset from GP, while Ferdous et al. also observed non-canonical cleavage in an independent Pallas dataset. Moreover, in a study performed by Curaba et al. (2012) 96 target genes of GP for miRNAs involved in seed development and germination were identified by degradome sequencing and only 16 targets were cleaved exclusively at the 10^th^-11^th^ nt site, while the other targets were sporadically cleaved with an offset (24) and 56 were cleaved in majority in a non-canonical site. Finally, Deng et al. (2015) identify in the barley cultivar Morex 65 target genes of 39 miRNAs, and for only 32% of the identified targets the canonical 10^th^-11^th^ nt cleavage product was the major degradome product. Together these studies highlight the challenges in the identification of cleavage sites of sRNAs in barley and cleavage sites of tRFs in plants. The absence of canonical cleavage products for tRFs does therefore not exclude the tRF-directed cleavage of *HvBAK1*, *HvEOL1* and *HvSERK2*.

We found that 22 *Fg*-sRNAs target *HvEOL1*, a putative negative regulator of ET biosynthesis in barley. In *Arabidopsis thaliana* the EOL1 homolog *At*ETO1 acts together with *At*EOL1 and *At*ETO1-like 2 (EOL2) in directing the ubiquitination and subsequent degradation of type-2 1-aminocyclopropane-1-carboxylate synthase (ACS) proteins (e.g. ET overproducer 2 (ETO2)) (Christians et al. 2009; Yoshida et al. 2006). ET is a gaseous plant hormone that plays an important role in regulating plant growth and development, and is critical for pathogen interaction and abiotic stresses (Abeles et al. 1992). Generally, ET acts synergistically with jasmonate (JA) in the defence response against necrotrophic pathogens and this ET/JA response has antagonistic effects on salicylic acid (SA) signalling against biotrophic pathogens. Yet in low amounts JA and SA act synergistically (Glazebrook 2005; Li et al. 2019). Therefore, controlling both ET biosynthesis and ET signalling is crucial for plants. Towards this, plants have evolved complex mechanisms that allow tight regulation of ET pathways e.g. at the level of (i) ET production mainly by regulating ACS gene family members, (ii) ET perception through constitutive triple response 1 (CTR1)-mediated inhibition of positive regulator ET insensitive 2 (EIN2) (Kieber et al. 1993, Alonso et al. 1999), and (iii) expression of ET-responsive TFs (e.g. ET response factor 1 (ERF1)) via EBF-mediated degradation of ET insensitive 3 (EIN3) (Potuschak et al. 2003) (Fig. S4). According to the anticipated role of ET in the plant response to necrotrophic pathogens, such as *Fg*, targeting negative regulators of ET synthesis such as *HvEOL1* would be detrimental to *Fg* colonization. Of note, our findings are consistent with previous results demonstrating that *Fg* exploits ET signalling to enhance colonization of Arabidopsis, wheat and barley (Chen et al. 2009), supposedly through an increase in DON-induced cell death through ET. These findings further challenge the role of ET in defence against necrotrophic pathogens. Strikingly, the authors showed that in Arabidopsis ET overproducing mutants (ETO1 and ETO2) and a negative regulator of ET signalling (CTR1) are more susceptible to *Fg*, while *At* mutants in ET perception (ETR1) and signalling (EIN2 and EIN3) are resistant. These findings were confirmed by the direct application of ET during the infection of wheat and barley, which lead to increased susceptibility to *Fg*. Based on these findings, we suggest that negative regulators of ET are efficient targets for sRNA-directed manipulation of host immunity by *Fg*.

The bacterial pathogen *Pseudomonas syringae* secretes two effector molecules, AvrPto and AvrPtoB, into host plants. These effectors interact with the receptor-like kinase BRI1-associated receptor kinase 1 (BAK1), also known as SERK3, thereby preventing the recognition of various MAMPs through the association of BAK1 with pattern recognition receptors (PRRs) such as flagellin-sensitive 2 (FLS2) and Ef-Tu receptor (EFR) (Shan et al. 2008). We observed *FgDCL*-dependent silencing of the cereal BAK1 homologs *HvBAK1*, *HvSERK2* and *BdSERK2*. While these genes have a higher similarity to *AtSERK2* than to *AtSERK3* (*AtBAK1*), they still are among the closest homologs to *AtBAK1* found in cereals (Fig. S9). It is tempting to speculate that further experiments will uncover additional hubs that are targeted both by protein and sRNA effectors.

## Conclusion

Our data show that in the necrotrophic ascomycete *Fusarium graminearum* gene silencing by RNAi shapes its ability to cause disease, which is consistent with earlier results on the significance of the RNAi machinery in *Fg* (Gaffar et al. 2019; Son et al. 2017). Pathogenicity relies on DICER-like (DCL)-dependent sRNAs that were identified as potential candidates for fungal effectors targeting defence genes in two Poaceae hosts, barley and *Brachypodium*. We identified *Fg*-sRNAs with sequence homology to host genes that were down-regulated by *Fg* during plant colonisation, while they were expressed above their level in healthy plants after infection with a DCL dKO mutant. In PH1-*dcl1/2 vs*. PH1 the strength of target gene accumulation correlated with the abundance of the corresponding *Fg*-sRNA. Our data hint to the possibility that three DCL-dependent tRFs with sequence homology to immunity-related *Ethylene overproducer 1-like 1* (*HvEOL1)* and three Poaceae orthologues of *Arabidopsis thaliana BRI1-associated receptor kinase 1* (*HvBAK1, HvSERK2* and *BdSERK2*) contribute to fungal virulence via targeted gene silencing.

## Experimental procedures

### Plants, fungi and plant infection

*Fusarium graminearum* (*Fg*) strain PH1, the double knock-out (dKO) PH1-*dcl1/2* (Dr. Martin Urban, Rothamsted Research, England), strain IFA65 (IFA, Department for Agrobiotechnology, Tulln, Austria) and single mutants IFA65-*dcl1* and IFA65-*dcl2* (Gaffar et al. 2019) were cultured on synthetic nutrient poor agar (SNA). Preparation of fungal inoculum was performed as described (Koch et al. 2013). *Arabidopsis thaliana* ecotype Col-0 and *Atago1-27* (Morel et al. 2002; Polymorphism:3510706481) were grown in 8 h photoperiod at 22°C with 60% relative humidity in a soil - sand mixture (4:1) (Fruhstorfer Type T, Hawita, Germany). For infection, 15 rosette leaves were detached and transferred in square Petri plates containing 1% water-agar. Drop-inoculation of Arabidopsis leaves was done with 5 µl of a suspension of 5 × 10^4^ *Fg* conidia ml^−1^ at two spots per leaf. Infection strength was recorded as infection area (size of chlorotic lesions relative to total leaf area) using the ImageJ software (https://imagej.nih.gov/ij/).

For infection of barley (*Hordeum vulgare* cv. Golden Promise, GP) and *Brachypodium distachyon* (Bd21-3), plants were grown in a 16 h photoperiod at 20°C/18°C day/night and 60% relative humidity in soil (Fruhstorfer Type LD80, Hawita). Ten detached second leaves were transferred into square Petri plates containing 1% water-agar. GP leaves were drop-inoculated with 3 µl of 1.5 x 10^5^ conidia ml^-1^ conidia suspension. Bd21-3 leaves were drop-inoculated on two spots with 10 µl of 1 x 10^4^ conidia ml^-1^ conidia suspension. Infection strength was measured with the PlantCV v2 software package (https://plantcv.danforthcenter.org/) by training a machine learning algorithm to recognize necrotic lesions. For gene expression analysis, a suspension of 5 x 10^4^ *Fg* conidia ml^-1^ was used and leaves were either inoculated on 3 spots with 20 µl (barley) or on 2 spots with 10 µl (Brachypodium), respectively and experiments were evaluated 5 days post inoculation (dpi).

### Fungal transcript analysis

Gene expression analysis was performed using reverse transcription quantitative PCR (RT-qPCR). RNA extraction was performed with GENEzol reagent (Geneaid) following the manufacturer’s instructions. Freshly extracted mRNA was used for cDNA synthesis using qScript^TM^ cDNA kit (Quantabio). For RT-qPCR, 10 ng of cDNA was used as template in the QuantStudio 5 Real-Time PCR system (Applied Biosystems). Amplifications were performed with 5 μl of SYBR^®^ green JumpStart Taq ReadyMix (Sigma-Aldrich) with 5 pmol oligonucleotides. Each sample had three technical repetitions. After an initial activation step at 95°C for 5 min, 40 cycles (95°C for 30 sec, 60°C for 30 sec, 72°C for 30 sec) were performed followed by a melt curve analysis (60°C-95°C, 0.075°C/s). Ct values were determined with the QuantStudio design and analysis software supplied with the instrument. Transcript levels were determined via the 2^-ΔΔCt^ method (Livak & Schmittgen 2001) by normalizing the amount of target transcript to the amount of the reference transcript *Elongation factor 1-alpha* (*EF1-a*, FGSG_08811) gene (Tab. S1).

### Plant transcript analysis

Leaves were shock frozen at 5 dpi and RT-qPCR was performed as described above. Reference genes were *Ubiquitin-40S ribosomal protein S27a-3* (HORVU1Hr1G023660) for GP and *Ubi4* (Bradi3g04730) for Bd21-3 according to Chambers et al. (2012) (Tab. S1). Primers were designed using Primer3 v2.4.0 (Untergasser et al. 2012).

### Spray application of dsRNA

Second leaves of 2 to 3-week-old GP were detached and transferred to square Petri plates containing 1% water agar. dsRNA was diluted in 500 μl water to a final concentration of 20 ng μl^-1^. As control, Tris-EDTA (TE) buffer was diluted in 500 μl water corresponding to the amount used for dilution of the dsRNA. Typical RNA concentration after elution was 500 ng/μl, with 400 μM Tris-HCL and 40 μM EDTA in the final dilution. Each plate containing 10 detached leaves was evenly sprayed with either dsRNAs or TE buffer with 500 µl, and subsequently kept at room temperature (Koch et al. 2016). Two days after spraying, leaves were drop-inoculated with three 20 µl drops of *Fg* suspension containing 5 × 10^4^ conidia ml^−1^. After inoculation, plates were closed and incubated for five days at room temperature.

### Target prediction for sRNAs

RNA was purified and enriched for sRNAs from fungal axenic culture using the mirVana miRNA Isolation Kit (Life Technologies). Indexed sRNA libraries were constructed from these sRNA fractions with the NEBNext Multiplex Small RNA Library Prep Set for Illumina (New England Biolabs) according to the manufacturer’s instructions. Reads were trimmed with the cutadapt tool v2.1 (Martin 2011) by removing adapters and retaining reads with a length of 21-24 nt and quality checked with the fastQC tool v0.11.9 (http://www.bioinformatics.babraham.ac.uk/projects/fastqc/). For Fig. S1 reads were aligned to the *Fg* reference genome (GCF_000240135.3_ASM24013v3) with bowtie2 (Langmead & Salzberg 2012) following a sensitive alignment policy (-D 100, -R 10, -L 19). The aligned reads were assigned to the additional attribute “gene_biotype” with htseq-count (Anders et al. 2015) according to the latest assembly (ftp://ftp.ensemblgenomes.org/pub/release-44/fungi/gff3/fungi_ascomycota3_collection/fusarium_graminearum_gca_000240135).

Remaining reads were collapsed with the fastx toolkit v0.0.14 (Hannon 2010) and reads with at least 400 reads were targeted against the IBSC_PGSB_v2 cDNA annotation with the plant miRNA target prediction algorithm TAPIR (Bonnet et al. 2010), following the optimized parameters according to Srivastava et al. (2014). The results of the target prediction were further analysed with RStudio (RStudio Team 2016) and the package biomaRt (Durinck et al. 2005) to find targets associated with stress and immunity associated Gene ontology (GO) terms in the database “plants_mart” from plants.ensembl.org hosted by the EBI (European Bioinformatics Institute) and the Wellcome Trust Sanger Institute. The same method was used for the identification of target genes in *B. distachyon* (GCA_000005505.4) and *A. thaliana* (Araport11).

### Stemloop-RT-qPCR of sRNAs

RNA was extracted and genomic DNA was digested as described above. The sequences of sRNAs found in axenic fungal culture were used to design specific stem loop (SL) primers matching the sRNA over 6 nt at the 3’end. For the primer design, the tool of Adhikari et al. (2013) was used. SL-primers were diluted to 10 pM and folded in a cycler (95°C for 15 min, 90°C 5 min, 85°C 5 min, 80°C 5 min, 75°C 1 h, 68°C 1 h, 65°C 1 h, 62°C 1 h, 60°C 3 h). These primers were used for cDNA synthesis (Thermo Scientific RevertAid RT Reverse Transcription Kit) according to manufacturer’s instruction with an annealing step at 16°C instead of 25°C and were used in multiplex to target respective fungal sRNAs and barley miRNAs *Hvu*-mir159 and *Hvu*-mir168 as references. To obtain amplification efficiencies, a mix from all RNAs was diluted in a four step dilution series with a factor of ten and reverse transcribed. Reactions were set up with the highest concentration of 15 ng µl^-1^ and the lowest of 15 pg µl^-1^ cDNA. All sRNA amplifications showed an efficiency of 80-82% and an R² between 1 and 0.997 except for Fg-sRNA-6717 with an efficiency of 66.4%. For RT-qPCR, 1.5 µl of 3 ng µl^-1^ cDNA was used as template in the QuantStudio 5 Real-Time PCR system (Applied Biosystems). Amplifications were performed with 5 μl of SYBR^®^ green JumpStart Taq ReadyMix (Sigma-Aldrich) with 1.5 pmol or 3 pmol oligonucleotides. Each sample had three technical repetitions. As forward primer the unused nucleotides of the remaining sequence of the sRNA were used, which were extended to achieve optimal melting temperature, and as reverse primer the universal stem loop primer developed by Chen et al. (2005) was used. Relative abundance of the sRNAs was calculated with the ΔΔct-method with incorporation of amplification efficiencies. sRNAs were normalized against the reference miRNAs Hvu-mir-159a and 168-5p and after this against the fungal biomass measured as *EF1-α* agains*t HvUBQ* (*HORVU1Hr1G023660*).

### Statistics

To assess the differential expression of genes via RT-qPCRs the Δct values were compared via a one or two sided paired Students *t*-test. Disease symptoms were either compared via Students *t*-test if the data showed a normal distribution in Shapiro-Wilk test or via a Wilcoxon rank sum test.

### RLM-RACE

RNA from GP barley infected with *Fg*-IFA65 at 5 dpi and an uninfected control was extracted with the Isolate II plant miRNA kit (Bioline). 1 µg of RNA (>200 nt) of infected, uninfected and a mix of both samples for a –RT-control were assembled. 1 µl of the 5’RACE Adapter [0.3 µg/µl], 1 µl of the 10x Reaction Buffer, 1 µl of 1mg/µl BSA, 0.5 µl of T4 RNA Ligase [10U/µl] (Thermo Scientific) and DEPC-treated water up to 10 µl were prepared and incubated at 37°C for 60 min. Subsequently, the whole reaction was used for reverse transcription (RevertAid Reverse Transcriptase, Thermo Scientific). 10 µl ligation reaction, 1 µl Random Hexamer [100pmol/µl], 4 µl 5x Reaction Buffer, 0.5 µl RiboLock RNase Inhibitor (Thermo Scientific), 2 µl dNTP Mix [10 mM] and 1 µl RevertAid Reverse Transcriptase (or water (–RT control)) and 1.5 µl water were mixed and run for 10 min at 25°C, 60 min at 42°C and 10 min at 70°C. Then, a nested hot-start touch-down PCR for each target gene was performed. The primer sequences for the outer (first) and inner (second) PCR are shown in Tab. S1. 5 µl of 10x Buffer B, 1 µl of a dNTP Mix [10 mM], 2 µl MgCl_2_ [25 mM], 1µl Adapter specific Primer [10 pmol µl^-1^] and 1 µl gene specific primer (GSP) [10 pmol µl^-1^], 0.6 µl DCS DNA Polymerase (DNA Cloning Service) [5 U/µl] and 2 µl cDNA or outer PCR reaction and 37.4 µl water were mixed and run at 95°C for 5 min, (95°for 30 s, 68°C-0.5°C/cycle for 30 s, 72°C for 30 s)*15, (95°C for 30 s, 60°C for 30 s, 72°C for 30 s)*18 and 72°C for 5 min. PCR products were evaluated in a 1.5% agarose gel and bands of the expected size, which were present in the infected but not uninfected samples, were excised. Products were cleaned with the Wizard SV Gel and PCR Clean-Up System (Promega) and cloned with the pGEM-T easy Vector Systems (Promega). For each band, five clones were picked for sequencing. Plasmids from O/N cultures were extracted with the Monarch Plasmid Miniprep Kit (New England Biolabs) and sent for sequencing to LGC genomics.

### Analysis of target genes and targeting sRNAs

After the initial target prediction an additional target prediction for the newly released cultivar specific genome (GCA_902500625) of barley cv. Golden Promise (GP) was conducted. Adapters were removed and reads were collapsed as described before for the target prediction. All sRNA sequences were read with SeqinR (v3.6-1; Charif & Lobry 2007) and stored in a list of SeqFastadna objects. To identify the homologous genes to the already identified targets in GP, the cDNA library was blasted with the command-line blast application (Nucleotide-Nucleotide BLAST 2.6.0+) (Camacho et al. 2009) against the identified target sequences from the IBSC_PGSB_v2 cDNA library with percent identity of 90 and a query coverage of 55% as cut-off values. All sRNAs with at least two reads were written to a file in chunks of 2000 each and ran against each individual target gene with TAPIR via the system2 function in R (R Core Team 2019) in the RStudio software. Results were collected, stored in a data.frame, and further analysed with R. sRNAs identified to target a gene of interest (GOI) were written to a fasta file with SeqinR and blasted against the rRNAs from the assemblies GCA_900044135.1 (*Fg*-PH1), GCA_000240135.3 (*Fg*-PH1) and the Fusarium rRNAs from the RNAcentral fungal ncRNA dataset (ftp://ftp.ebi.ac.uk/pub/databases/RNAcentral/current_release/sequences/by-database/ensembl_fungi.fasta (12/Sep/2020)) with the options wordsize=4, perc_identity=95, qcov_hsp_perc=95. All sRNAs matching rRNAs were removed. Thereafter, sRNAs were compared to the *Fg* assemblies GCA_900044135.1 (*Fg*-PH1) and GCA_000240135.3 (*Fg*-PH1) with the same blast strategy and only perfectly matching sRNAs were retained.

To derive the relative expression of a GOI between two samples the following formula is used.

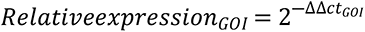

We further defined the ΔΔΔct value as the difference between the ΔΔct values for a GOI in PH1 and PH1-*dcl1/2-*infected samples.

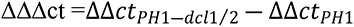

This enables the calculation of the re-accumulation between the two samples as follows.

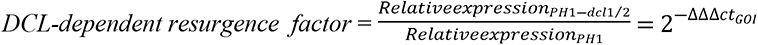

The sum of all reads and the corresponding ΔΔΔct-value were plotted with ggplot2 (Wickham 2016) and a linear regression was added to the plot. To allow a log2-transformation of the plots genes with zero targeting reads were set to one targeting read. The plots were arranged using ggpubr v.0.4.0 (Kassambara 2017).

### GO enrichment analysis

Gene ontology (GO) enrichment analysis was performed via the AgriGO v.2.0 analysis toolkit (Tian 2017) with the standard parameters singular enrichment analysis (SEA).

### Phylogenetic analysis of SERK homologs

Homologs of *HvBAK1* and *HvSERK2* were searched in *At*, *Hv* and *Bd* with biomaRt v.2.40.5 (Durinck et al. 2005) and downloaded from the EMBL’s European Bioinformatics Institute plants genome page (plants.ensembl.org) in the plants_mart dataset hvulgare_eg_gene (Ensembl Plants Genes v. 50). For these homologs the CDS of all homologs within the respective datasets athaliana_eg_gene, hvulgare_eg_gene and bdistachyon_eg_gene were downloaded. The CDS were subsequently aligned with the muscle algorithm in MEGA7 (Kumar et al. 2016b) and a phylogenetic tree was constructed via a bootstrap method with 200 iterations.

## Acknowledgements

We thank the Salk Institute Genomic Analysis Laboratory for providing the sequence-indexed Arabidopsis TDNA insertion mutants. This work was supported by the Deutsche Forschungsgemeinschaft (Research Unit FOR5116) to KHK.

## Supporting Information

**Fig. S1:**
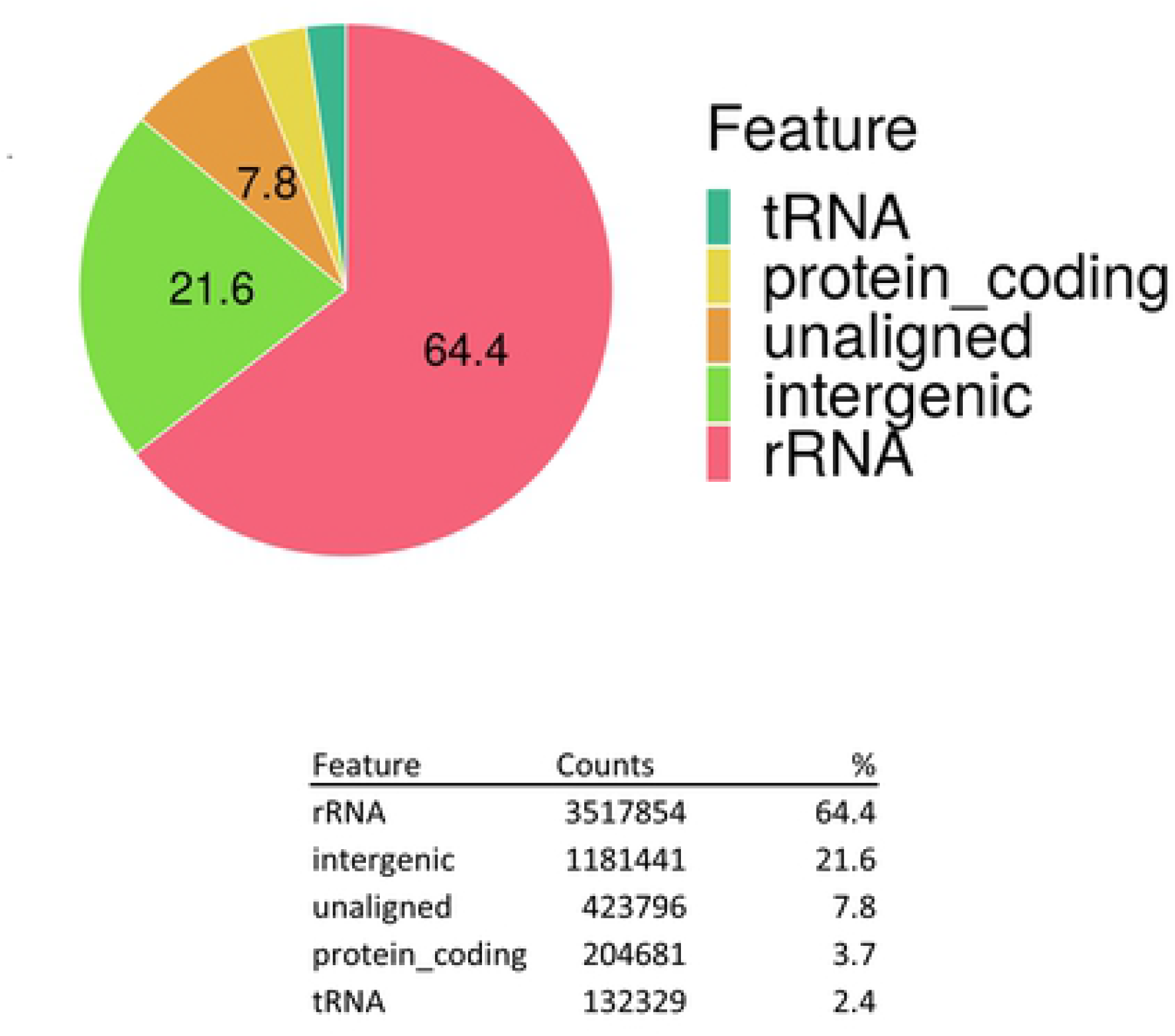
Feature mapping of Fg-sRNAs with a read length of 21-24 nt. Reads were trimmed as described earlier and aligned to the PH1 reference genome (GCF_000240135.3_ASM24013v3) with bowtie2 (Langmead & Salzberg 2012).

**Fig. S2:**
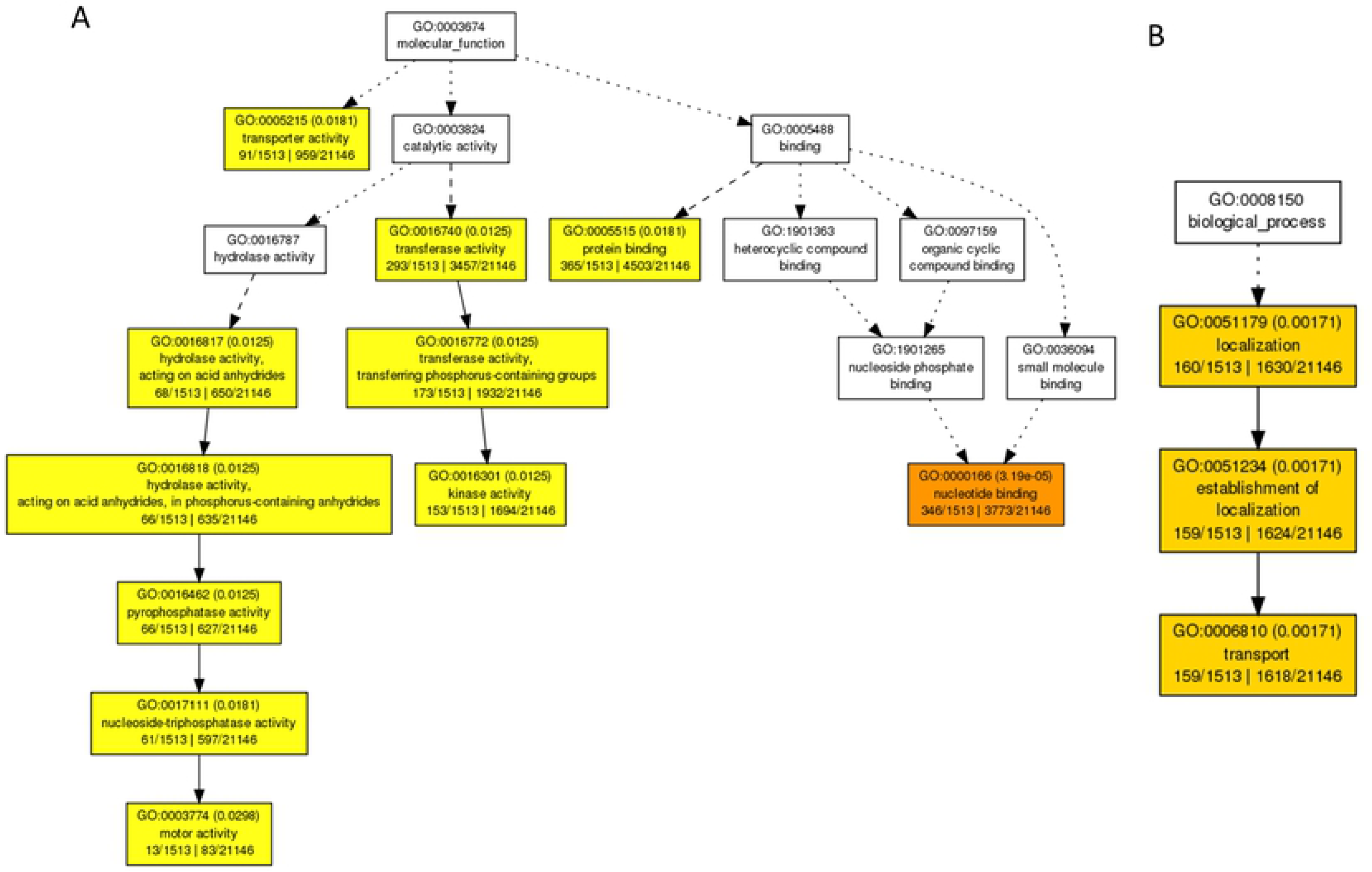
GO-enrichment analysis of all potential targets of Fg-sRNAs with more than 400 reads. The plot shows all significantly enriched GO-terms in the target gene set for (A) molecular function and (B) biological process. The analysis was done using agriGo v2.0. Each box contains information regarding one term. GO: indicates the GO accession, in brackets the p-value is stated (Fisher; Yekutieli (FDR)). After the bracket the GO-term description is written followed by the number of genes associated with said term 1. in the gene set and 2. In the background.

**Fig. S3:**
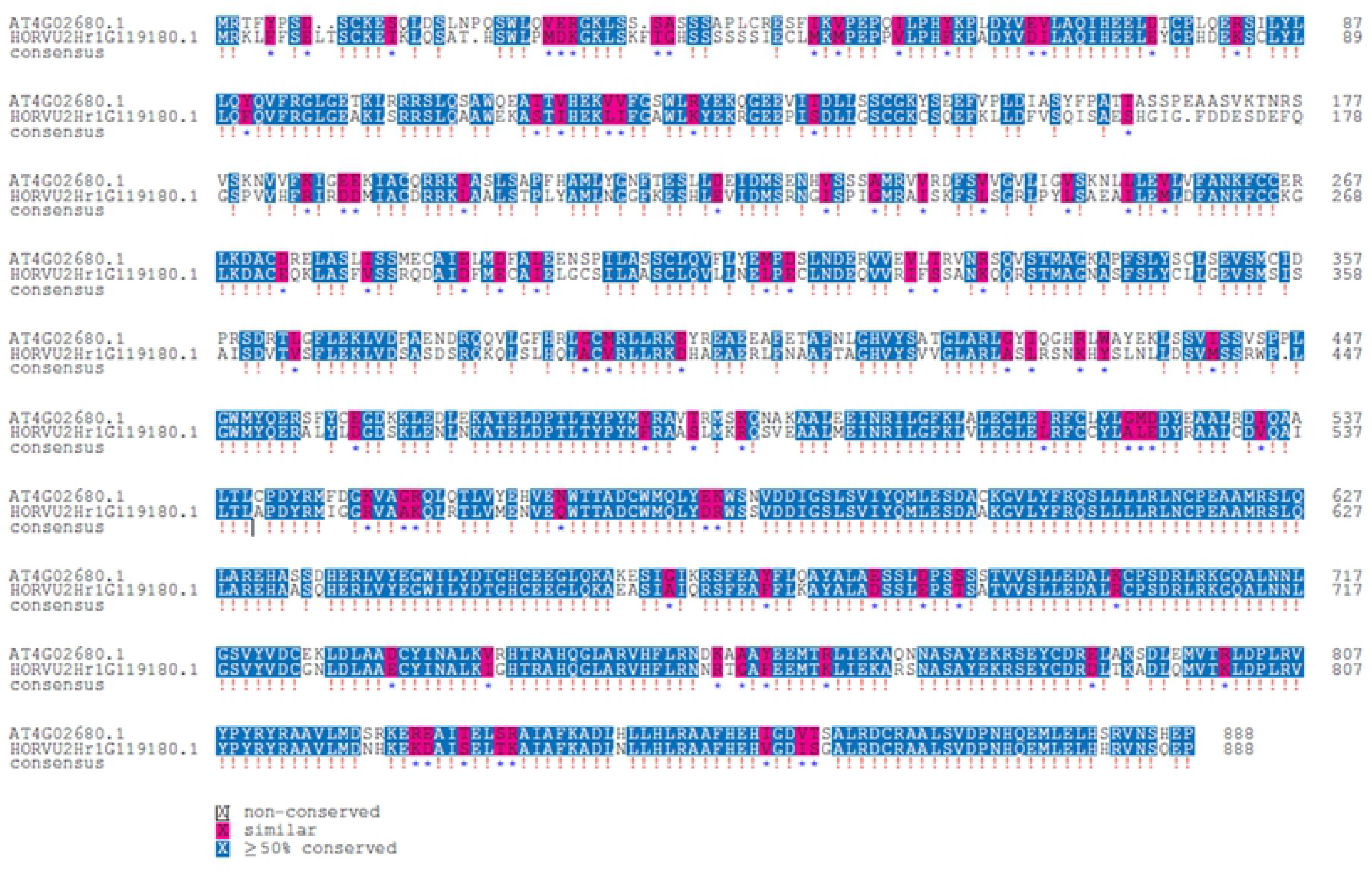
Alignment of AtETO1 and HvEOL1. Identical amino acids are marked blue and similar amino acids are marked red. The alignment and visualization was done with the msa package for R (Bodenhofer et al. 2015).

**Fig. S4:**
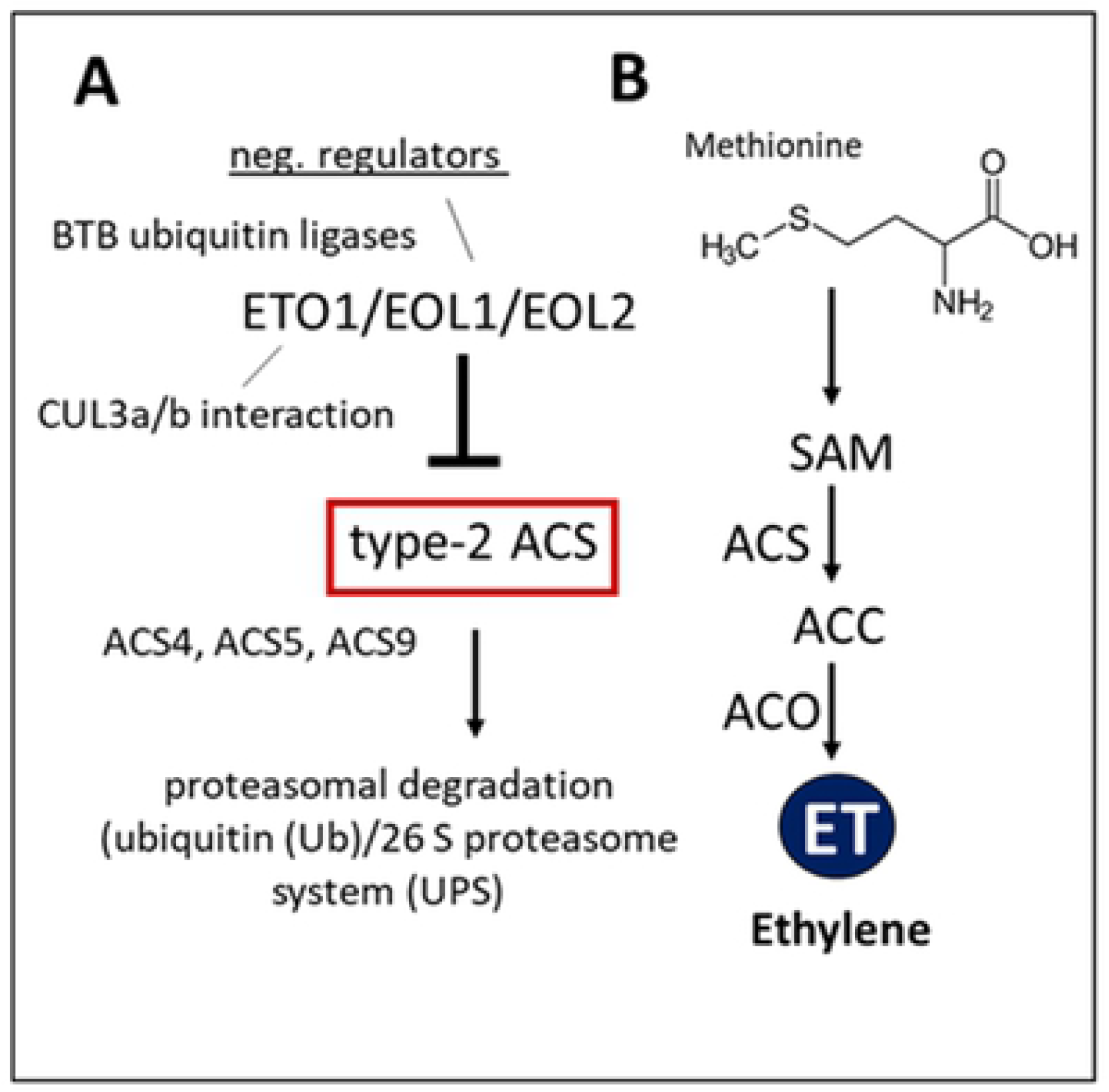
Regulation of ET synthesis in At. *At*ETO1 negatively regulates ethylene **(ET)** synthesis in *At*. *At*ETO1 acts together with *At*EOL1 and *At*ETO1-like 2 (EOL2) in directing the ubiquitination and subsequent degradation of type-2 1-aminocyclopropane-1-carboxylate synthase (ACS) proteins (e.g. ET overproducer 2 (ETO2)), which produce the direct precursor of ET.

**Fig S5:**
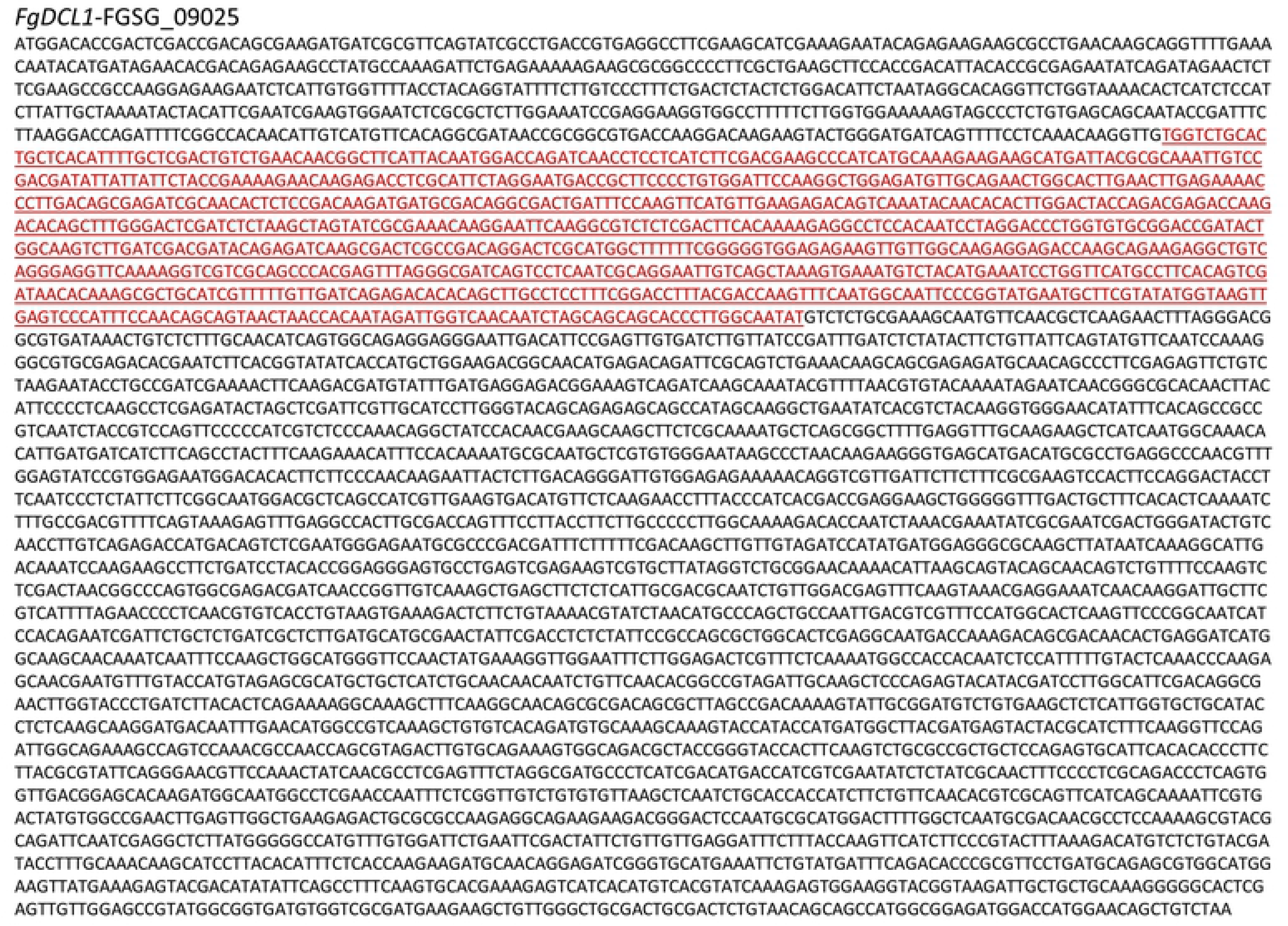
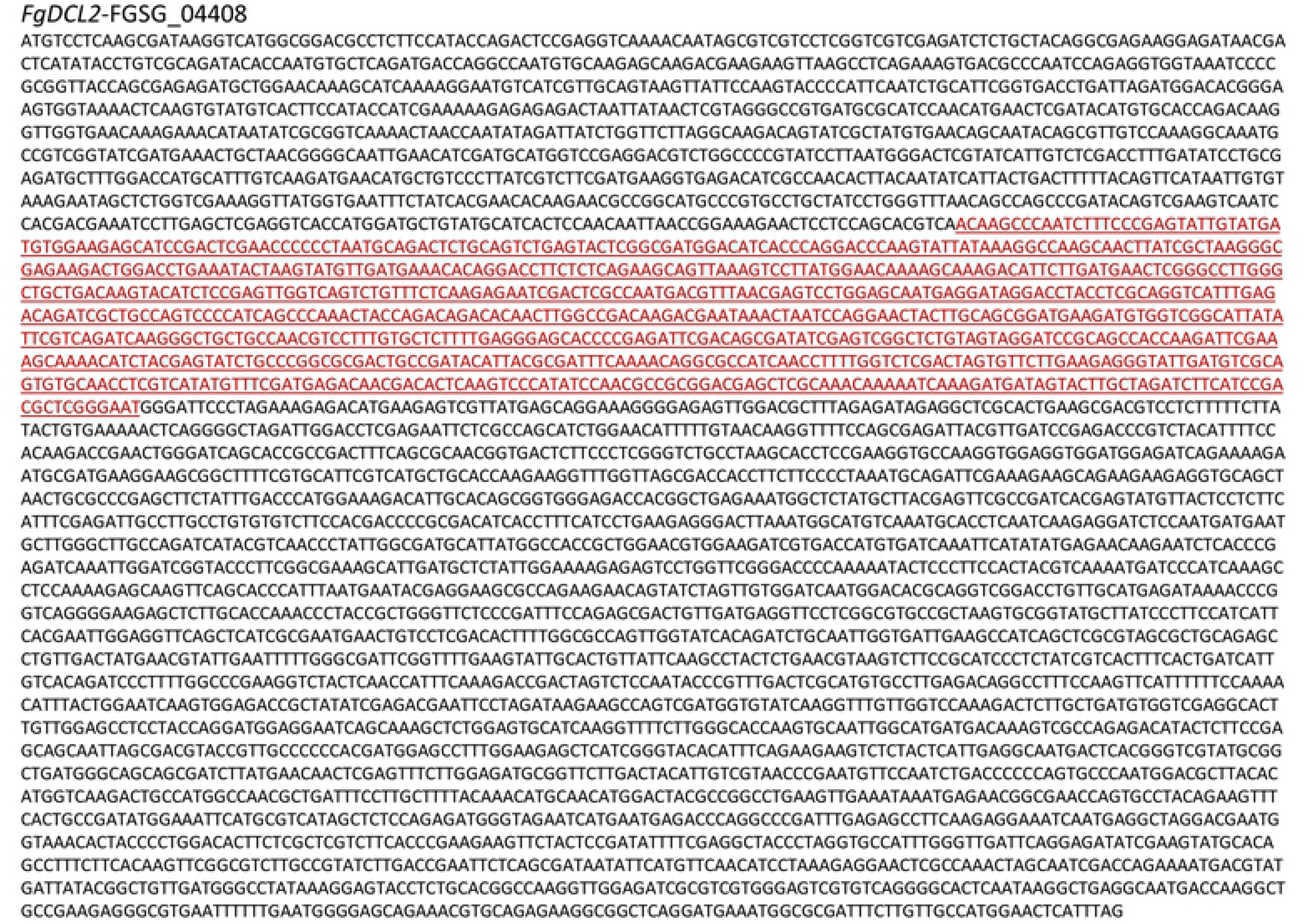
Sequences of dsRNA-dcl1/2. Coding Sequences (CDS) of the respective *FgDCL* genes with the sequences comprising the dsRNAs marked in red. **A.** *FgDCL1-*FGSG_09025 (912 nt long dsRNA-*Fg*DCL1). **B.** *FgDCL2-*FGSG_04408 (870 nt long dsRNA-*Fg*DCL2).

**Fig. S6:**
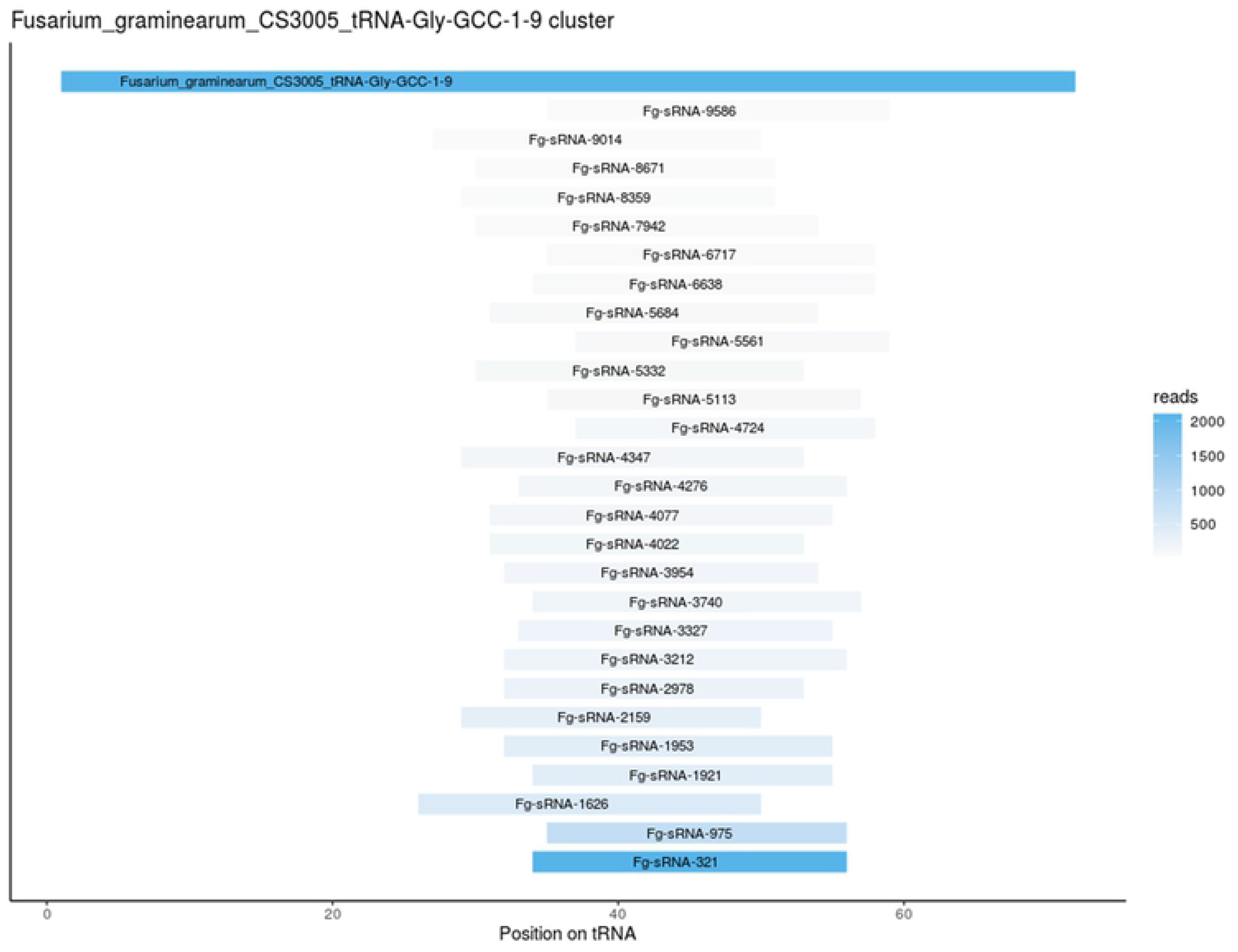
Position and read count of all tRFs from Fg-tRNA-Gly(GCC) Alignment position of all *Fg*-sRNAs from axenic culture with more than 50 reads perfectly matching the *Fg*-tRNA-Gly(GCC)-9 gene (Fusarium_graminearum_CS3005-tRNA-Gly-GCC-1-9) colored by read count.

**Fig. S7:**
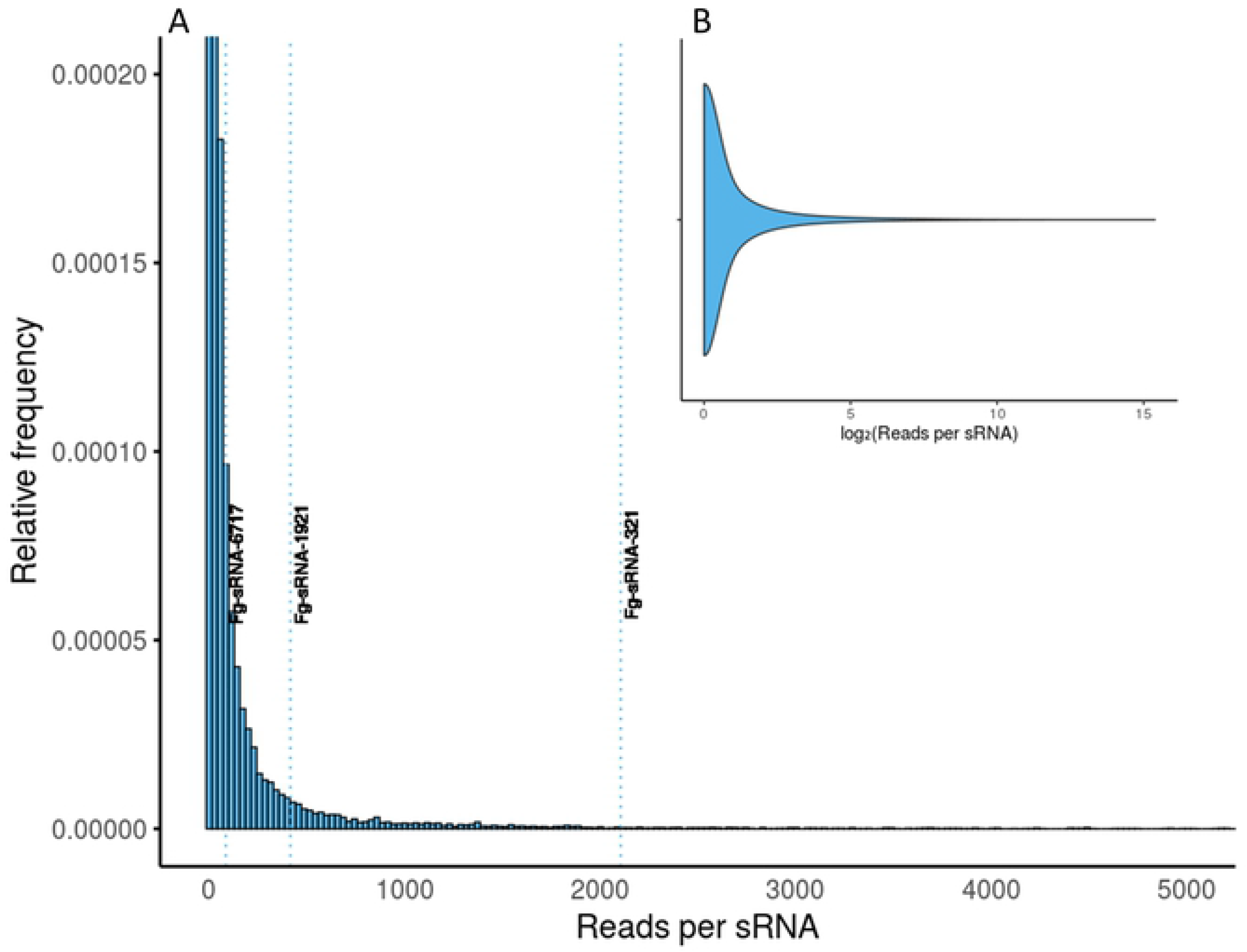
Abundance of unique Fg-sRNAs in axenic culture of IFA65. A: Histogram of the read count of every unique sRNA. The plot is truncated to make abundances recognizable. Most sRNAs have very low read counts and very few sRNAs have more reads than 3,000. Maximum read count per sRNA is 42,866. B: Violin plot of log2-transformed reads counts untruncated.

**Fig. S8:**
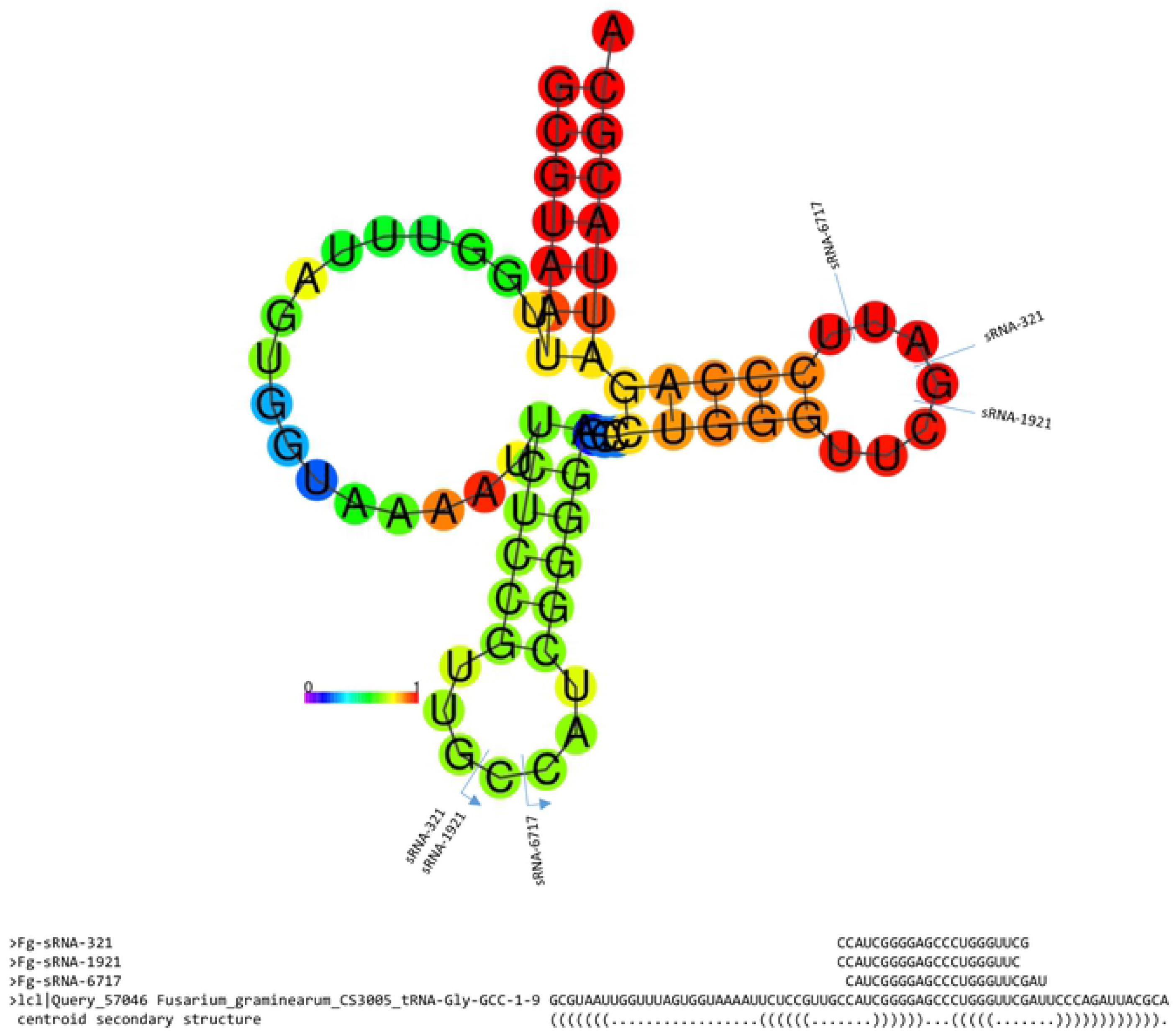
Origin of tRFs in Fg-tRNA-Gly(GCC) The centroid secondary structure of the *Fg*-tRNA-Gly(GCC) generated on the RNAfold web server (http://rna.tbi.univie.ac.at/cgi-bin/RNAWebSuite/RNAfold.cgi) with the origin and alignment of *Fg*-sRNA-321, *Fg*-sRNA-1921 and *Fg*-sRNA-6717. The colors of bases indicate the base pair probabilities.

**Fig. S9:**
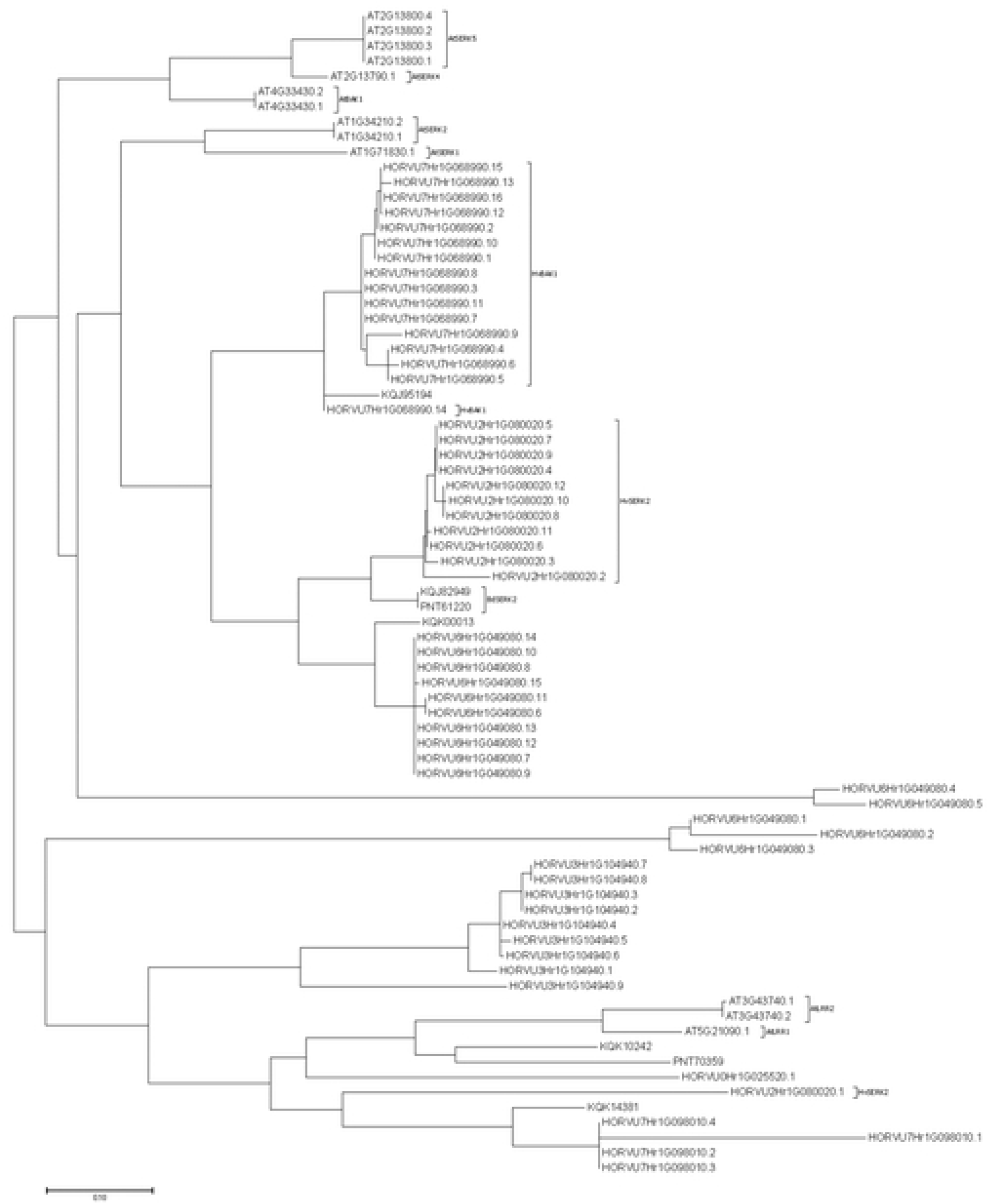
Molecular Phylogenetic analysis by Maximum Likelihood method. The evolutionary history was inferred by using the Maximum Likelihood method based on the General Time Reversible model (Nei & Kumar, 2000). The tree with the highest log likelihood (-25430.37) is shown. Initial tree(s) for the heuristic search were obtained automatically by applying Neighbor-Join and BioNJ algorithms to a matrix of pairwise distances estimated using the Maximum Composite Likelihood (MCL) approach, and then selecting the topology with superior log likelihood value. The tree is drawn to scale, with branch lengths measured in the number of substitutions per site. The analysis involved 77 nucleotide sequences. Codon positions included were 1st+2nd+3rd. There were a total of 2427 positions in the final dataset. Evolutionary analyses were conducted in MEGA7 (Kumar et al. 2016b).

**Tab. S1:**
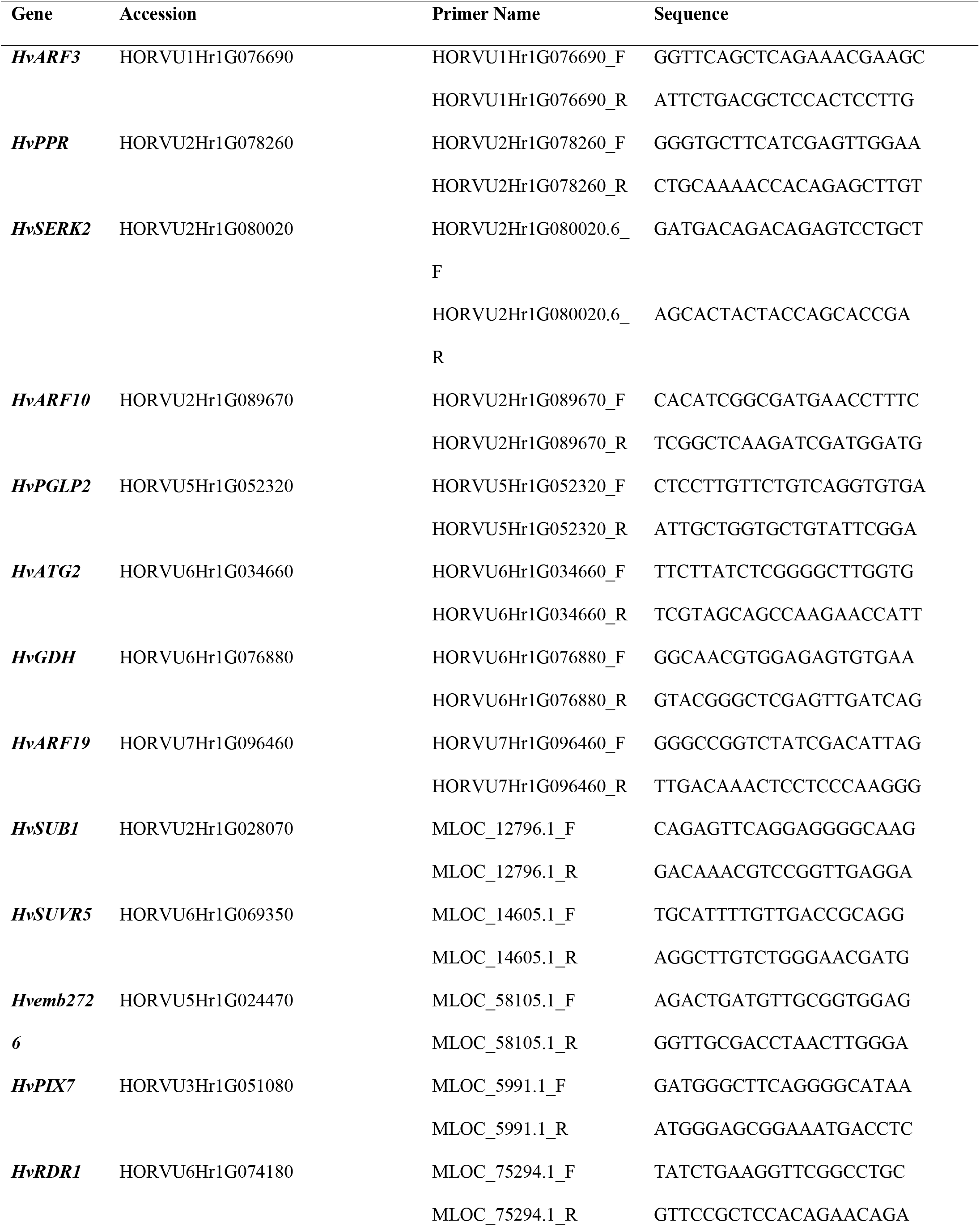
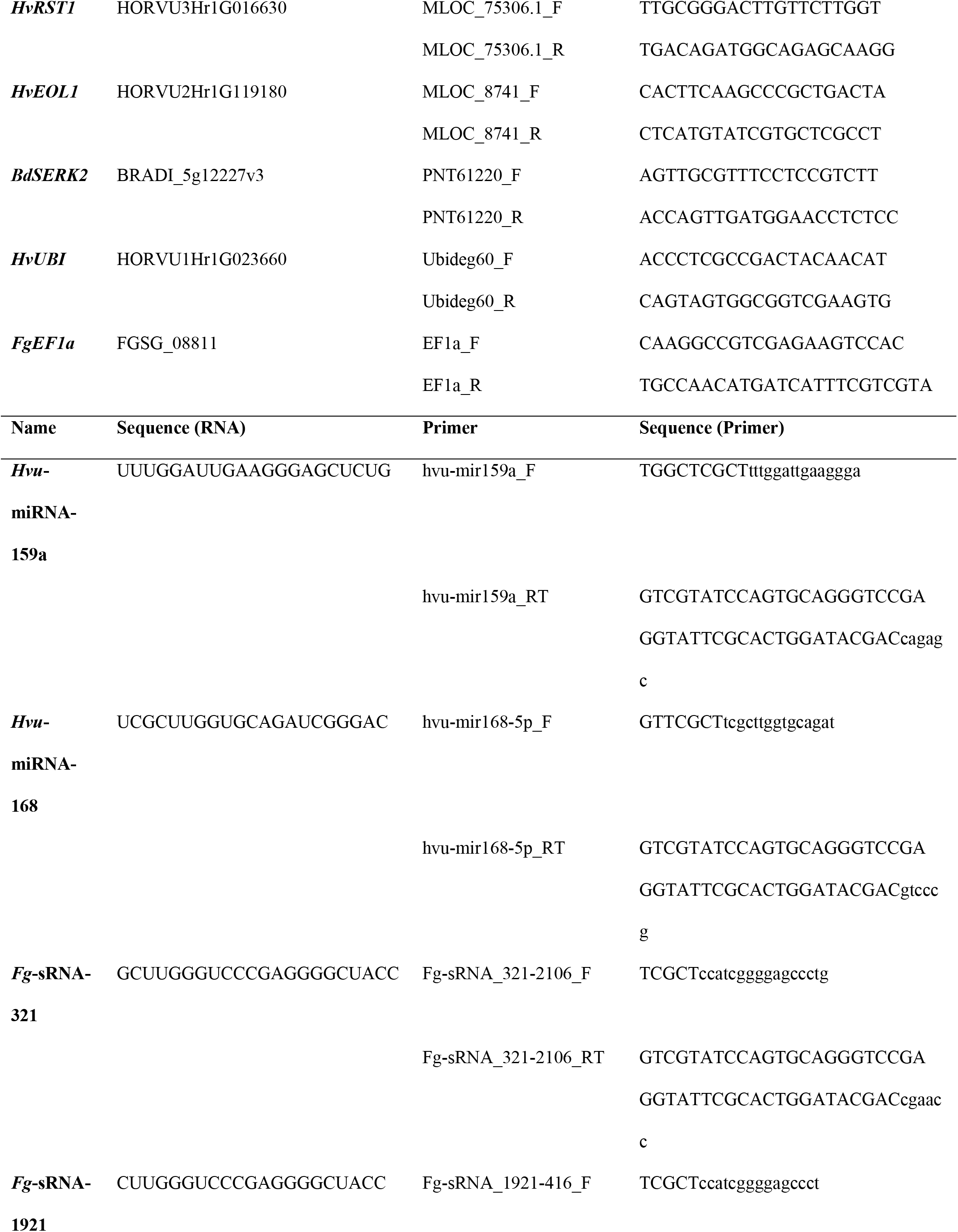
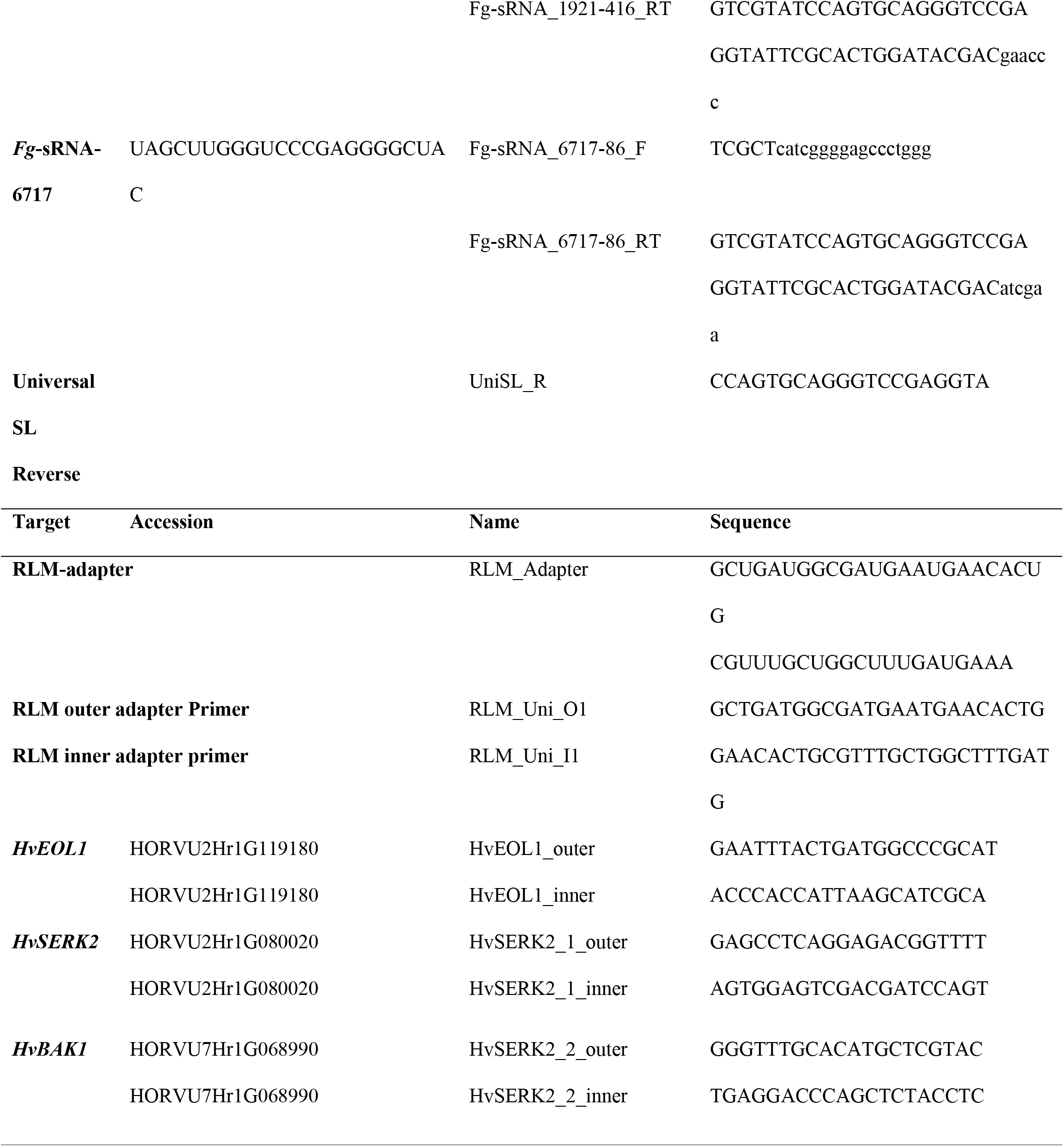
Primer sequences. Sequences and target accessions for all primers used in the study

**Tab. S2:** Target prediction results. Results of the target prediction with the TAPIR algorithm for all *Fg*-sRNAs with more than 400 reads

